# CD73 restrains mutant β-catenin oncogenic activity in endometrial carcinomas

**DOI:** 10.1101/2024.11.18.624183

**Authors:** Rebecca M. Hirsch, Sunthoshini Premsankar, Katherine C. Kurnit, Lilly F. Chiou, Emily M. Rabjohns, Hannah N. Lee, Russell R. Broaddus, Cyrus Vaziri, Jessica L. Bowser

## Abstract

Missense mutations in exon 3 of *CTNNB1*, the gene encoding β-catenin, are associated with poor outcomes in endometrial carcinomas (EC). Clinically, *CTNNB1* mutation status has been difficult to use as a predictive biomarker as β-catenin oncogenic activity is modified by other factors, and these determinants are unknown. Here we reveal that CD73 restrains the oncogenic activity of exon 3 β-catenin mutants, and its loss associates with recurrence. Using 7 patient-specific mutants, with genetic deletion or ectopic expression of CD73, we show that CD73 loss increases β-catenin-TCF/LEF transcriptional activity. In cells lacking CD73, membrane levels of mutant β-catenin decreased which corresponded with increased levels of nuclear and chromatin-bound mutant β-catenin. These results suggest CD73 sequesters mutant β-catenin to the membrane to limit its oncogenic activity. Adenosine A1 receptor deletion phenocopied increased β-catenin-TCF/LEF activity seen with *NT5E* deletion, suggesting that the effect of CD73 loss on mutant β-catenin is mediated via attenuation of adenosine receptor signaling. RNA-seq analyses revealed that *NT5E* deletion alone drives pro-tumor Wnt/β-catenin gene expression and, with CD73 loss, β-catenin mutants dysregulate zinc-finger and non-coding RNA gene expression. We identify CD73 as a novel regulator of oncogenic β-catenin and help explain variability in patient outcomes in *CTNNB1* mutant EC.

**Graphical Abstract.**
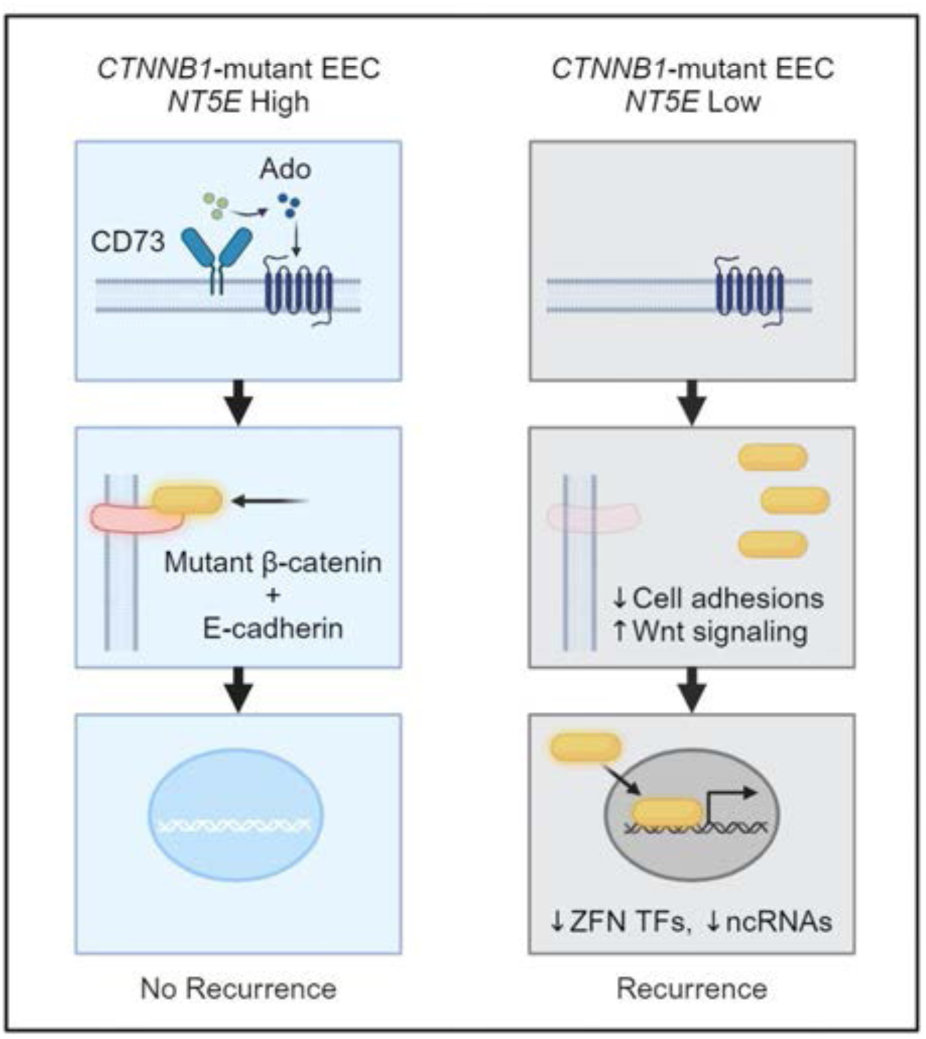

## Introduction

β-catenin, encoded by the gene *CTNNB1,* is an essential component of cell-cell adhesions^1–5^ and a transcriptional co-activator of Wnt signaling^6–10^. β-catenin is a crucial oncogene in various types of cancer and is often activated by somatic missense mutations in exon 3 of *CTNNB1*^11–15^. In endometrial carcinomas (EC), exon 3 *CTNNB1* mutations occur frequently (on average of ∼20-30%), especially in low grade, early stage endometrioid EC (EEC)^16–19^. Exon 3 mutations prevent β-catenin protein degradation, leading to its cytoplasmic accumulation and subsequent nuclear translocation and oncogenic transcriptional activity^13,14,20^. Several studies implicate *CTNNB1* mutations as oncogenic drivers in EC. For instance, *CTNNB1* mutations are seen as early as atypical hyperplasia and expression of a dominant stabilized *CTNNB1* exon 3 in murine studies results in endometrial hyperplasia^21–23^. Many more studies report that high expression of β-catenin or *CTNNB1* mutations are associated with recurrence, worse recurrence-free survival, or overall survival^16,17,24–28^ and that *CTNNB1* mutation is a greater risk factor for recurrence than other aggressive clinical features, such as myometrial invasion and lymphovascular space invasion (LVSI)^17^.

Despite a close association with recurrence, *CTNNB1* status has been challenging to utilize clinically for identifying patients at high risk for poor outcomes. *CTNNB1* mutation alone has poor sensitivity and specificity in predicting disease recurrence. Some patients with *CTNNB1* mutant tumors will have disease recurrence (∼30%), yet many others will never recur^28^. Additionally, studies assessing the nuclear localization of β- catenin by immunohistochemistry as a possible approach for predicting recurrence have shown that the percent of β-catenin nuclear expression in *CTNNB1* mutant EC^29^ and/or endometrial tumors with aberrant β-catenin expression^30^ is widely variable (for example, ∼5-60%^29^) with no clear correlation with outcomes. The variability in outcomes has suggested that β-catenin oncogenic activity is modified by other factors. To improve patient outcomes, it is critical to identify these determinates.

The purpose of this study was to examine whether CD73, a 5’-nucleotidase, is a critical factor controlling the oncogenic activity of exon 3 mutant β-catenin in EEC. In previous studies elucidating the role of CD73 in the pathogenesis of EC^31,32^, we found that CD73 localizes wildtype β-catenin to the membrane to prevent disease aggressiveness^31^. Unlike other cancers, CD73 is downregulated in EC^31,32^, and its loss is associated with worse overall survival^31^. CD73 limits EC cell invasion by activating adenosine A_1_ receptor (A_1_R) signaling, which redistributes E-cadherin and β-catenin to the cell membrane to protect epithelial integrity by reforming cell-cell adhesions^31^. In EEC, the majority of mutations in *CTNNB1* occur in exon 3 (88.7%; TCGA) in a stretch of 14 amino acids (codon 32 to 45)^16^. Exon 3 encodes a region on the N-terminus of β- catenin, which is separate from the region (armadillo domain repeats) where β-catenin binds with E-cadherin to localize to the membrane^33–36^. Based on our previous findings that CD73-A_1_R signaling redistributes wildtype β-catenin to the cell membrane, we hypothesized that CD73 limits the oncogenic activity of exon 3 mutant β-catenin by sequestering it to the membrane. Thus, CD73 serves as a molecular determinant of β- catenin oncogenic activity, whereby the presence or absence of CD73 expression in exon 3 *CTNNB1* mutant tumors helps explain the variability in patient outcomes.

Here we reveal that CD73 critically restrains the oncogenic activity of patient-specific exon 3 β-catenin mutants by sequestering mutant β-catenin to the cell membrane. We provide evidence for CD73 as a potential biomarker for predicting recurrence in patients with tumors with exon 3 *CTNNB1* mutation and uncover that CD73 loss alone in EC cells induces pro-tumor Wnt/β-catenin gene target expression, in addition to promoting novel β-catenin mutant-mediated changes in gene expression.

## Results

### Low *NT5E* expression associates with recurrence in endometrial cancer patients with exon 3 *CTNNB1* mutations

We first assessed whether CD73 was associated with disease recurrence in β-catenin mutant EC. *NT5E* expression was measured in *n* = 29 endometrial tumors verified by next-generation sequencing to have exon 3 *CTNNB1* mutations. Tumors were then stratified by patient recurrence or death (Supplemental Table 1). *NT5E* expression was significantly lower in patients with recurrence or who died of their disease compared to patients with no recurrence (Figure 1A). *NT5E* expression was not significantly different when the tumors were stratified by surgical stage (Figure 1B) or lymphovascular space invasion (LVSI) (Figure 1C), indicating that *NT5E* expression associates with recurrence. *NT5E* expression plotted for individual patients revealed the lower quartile contained most of the recurrences (Figure 1D). Of these patients, 86% had recurrence or death. Notably, the lower quartile value (0.00984 molecules of *NT5E*) is similar to values we previously reported to be associated with poor patient outcomes in a more diverse group of endometrial tumors^31^. Due to the small cohort size, we were not statistically powered for performing survival analyses. However, visible downward trends for progression-free survival and overall survival were observed in patients with *NT5E*-low tumors (Supplemental Figure 1).

**Figure 1.**
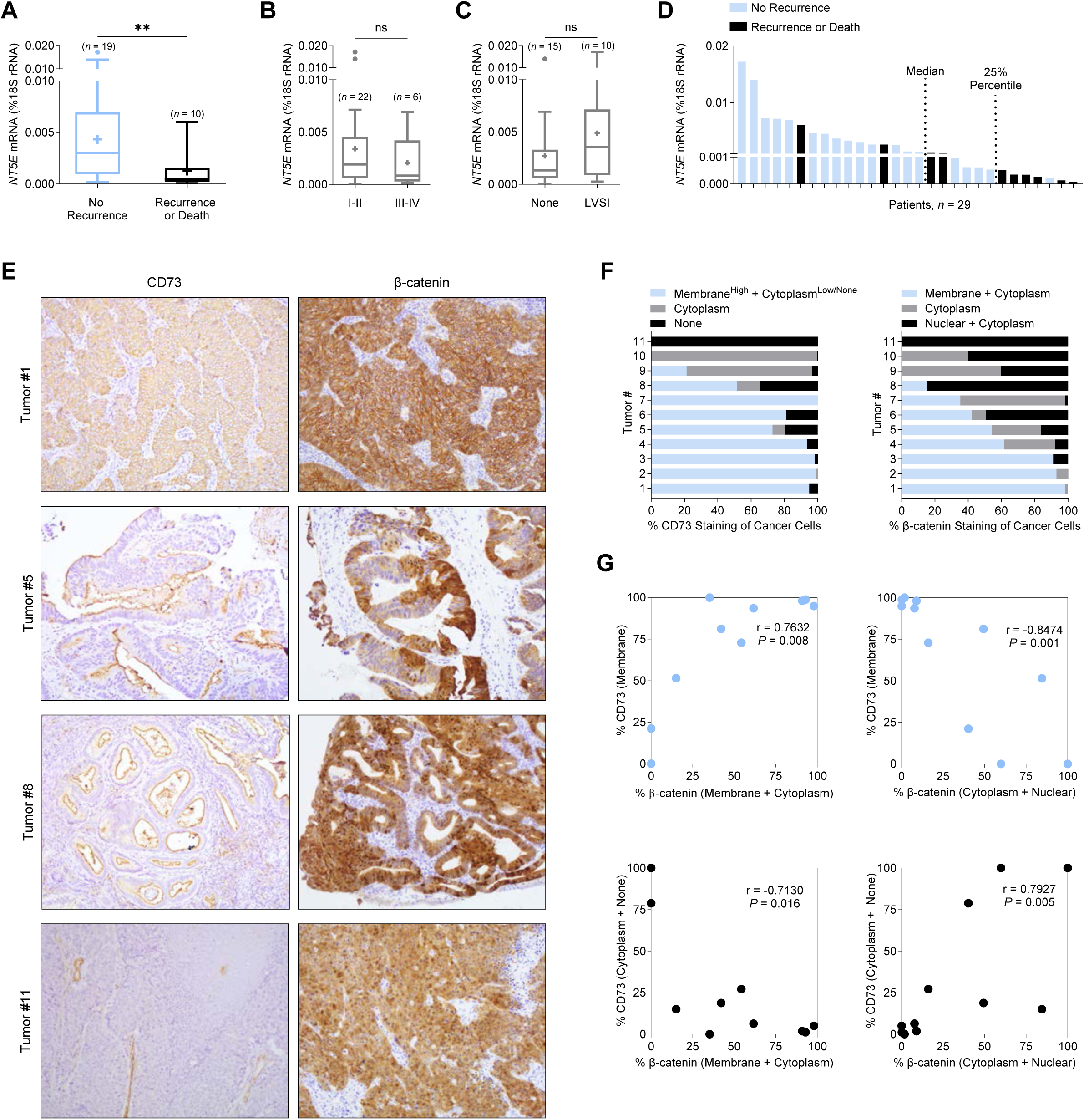
Loss of CD73 associates with poor patient outcomes and increased cytoplasmic/nuclear β-catenin in endometrial tumors with exon 3 *CTNNB1* mutations. (**A-D**) *NT5E* mRNA expression in exon 3 *CTNNB1* mutant endometrial tumors, stratified by (**A**) disease recurrence (*n* = 29), (**B**) International Federation of Gynecology and Obstetrics (FIGO) surgical stage (*n* = 28) and (**C**) lymphovascular space invasion (LVSI) (*n* = 25). FIGO surgical stage and LVSI information were not available for all 29 patients. Box blots represent the 25th–75th percentiles, bars are the 50th percentile, crosses are the mean values, and whiskers represent the 75th percentile plus and the 25th percentile minus 1.5 times the interquartile range. Values greater than these are plotted as individual circles. Data are presented as the molecules of *NT5E* transcript/molecules of 18S rRNA. (**D**) *NT5E* mRNA expression for *n* = 29 individual patient tumors, showing 6/7 patients recur with *NT5E* expression below the 25^th^ percentile. (**E**) Representative images of CD73 and β-catenin staining patterns for endometrioid endometrial carcinomas validated by next-generation sequencing to have an exon 3 *CTNNB1* (β-catenin) mutation. Tumor 1: membrane CD73 and β-catenin expression. Tumor 5: reduced membrane CD73 expression and membrane/cytoplasm/nuclear β-catenin expression. Tumor 8: minimal membrane CD73 expression and mostly cytoplasmic/nuclear β-catenin expression. Tumor 11: loss of CD73 expression and fully cytoplasmic/nuclear β-catenin expression. (**E**) Quantification of staining patterns for CD73 and β-catenin for *n* = 11 individual tumors. Data represents percent (%) staining pattern of cancer cells/total area of cancer cells. (**F**) Pearson correlations of data shown in **(E)**. **P < 0.01; Mann-Whitney test.

### Loss of CD73 associates with β-catenin nuclear localization in exon 3 *CTNNB1* mutant tumors

We found a strong correlation between CD73 expression and β-catenin localization in exon 3 *CTNNB1* mutant tumors, supporting our hypothesis that CD73 may control mutant β-catenin signaling. Immunohistochemistry studies showed CD73 loss was strongly correlated with cytoplasmic and nuclear β-catenin staining in EC cells in *CTNNB1*-mutant tumors (Figure 1E, Tumors #5, #8, and #11; Figure 1F and 1G). In contrast, CD73 membrane expression was strongly correlated with regions of the tumor with EC cells with membrane β-catenin staining (Figure 1E, Tumor #1; Figure 1F and 1G). Membrane expression of CD73 is indicative of cells maintaining epithelial integrity^31^. Based on these data, we further pursued *in vitro* experiments to test whether CD73 restrains the oncogenic activity of exon 3 mutant β-catenin, thus explaining the lower rates of recurrence in patients with ‘*NT5E* high’ tumors.

### Validation of the TOPFlash reporter assay for measuring mutant β-catenin activity in EC cells

Given that low *NT5E* expression was associated with disease recurrence, we sought to determine whether CD73 may control the oncogenic activity of mutant β-catenin in EC. β-catenin is a transcriptional co-factor and often complexes with proteins of the T-cell factor/lymphoid enhancer factor (TCF/LEF) family of transcription factors to activate the transcription of Wnt target genes^8–10,12,13,37^. Accordingly, we performed experiments testing a TOPFlash reporter as a readout of mutant β-catenin transcriptional activity in EC cells^12^. The TOPFlash construct contained 8 TCF/LEF binding sites upstream of a luciferase gene promoter^38^. HEC-1-A and Ishikawa cells are EC cell lines commonly used to model low grade EC and express high or low/no levels of CD73, respectively (Fig 2A-2B)^31^. We also used a multi-site exon 3 *CTNNB1 Xenopus* mutant (*Xenopus* β-catenin^ΔEX3^) to test the induction of TCF/LEF reporter activity^39^. *Xenopus CTNNB1* is 97% homologous to the human *CTNNB1* gene and 100% homologous in the exon 3 region (UniProt, P35222 and P26233). Our validation experiments showed both endogenous β-catenin and *Xenopus* β-catenin^ΔEX3^ induced TCF/LEF reporter activity in HEC-1-A and Ishikawa cells (Figure 2C, 2F), with *Xenopus* β-catenin^ΔEX3^ showing a greater induction (Figure 2C, 2F). TCF/LEF activity was not induced in cells transfected with FOPFlash, a construct with mutated TCF/LEF sites^38^ (Figure 2C), and was reduced by *CTNNB1* siRNA (Figure 2D-2E), demonstrating that the luciferase signals of the TOPFlash reporter were specific to β-catenin.

**Figure 2.**
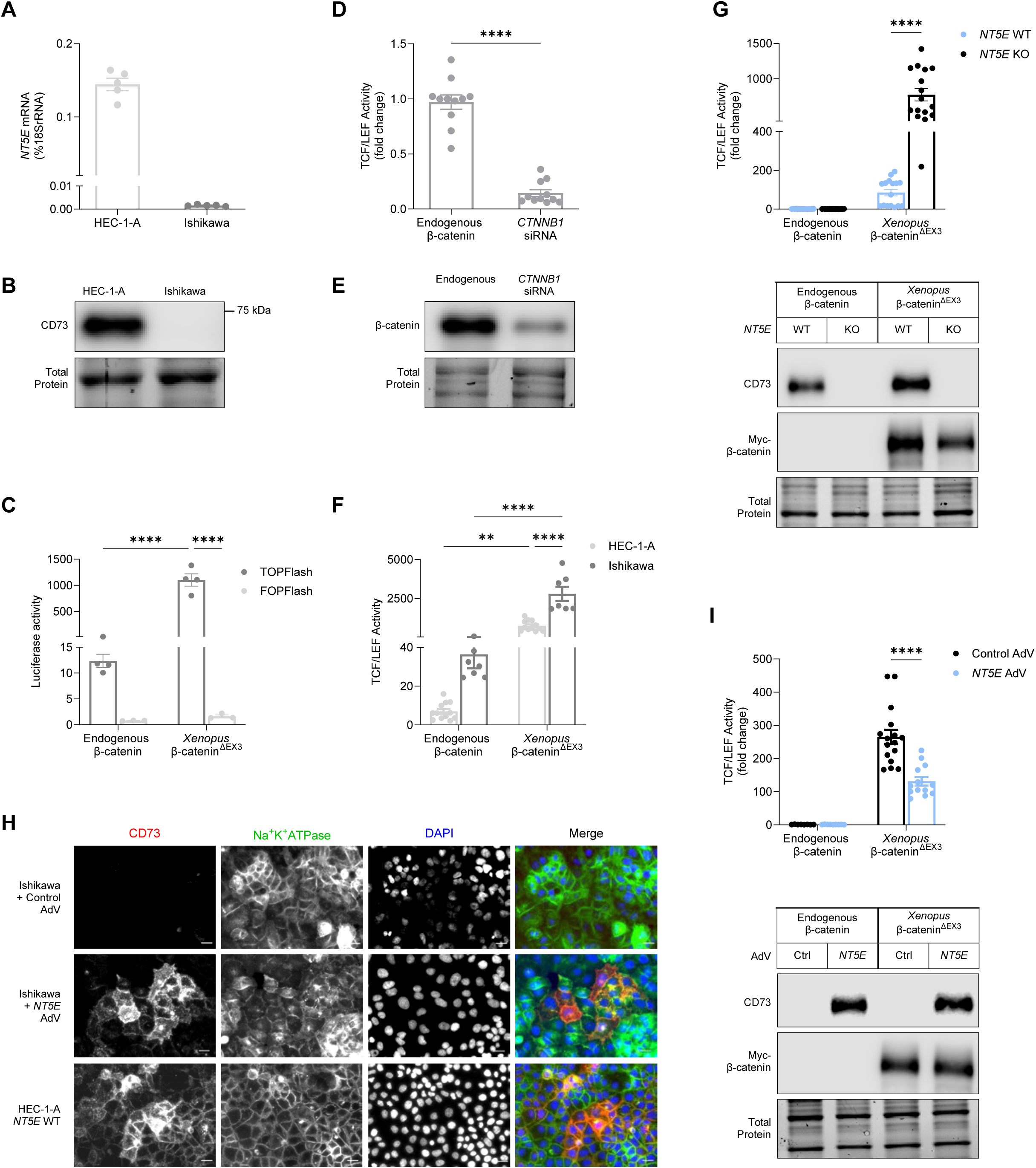
CD73 restrains the transcriptional activity of *Xenopus* exon 3 mutant β- catenin. (**A**) *NT5E* mRNA and (**B**) protein (CD73) expression in EC cell line models. (**C**) Validation of TCF/LEF-luciferase reporter activity using TOPFlash/FOPFlash plasmids for endogenous β-catenin and *Xenopus* exon 3 mutant β-catenin (β-catenin^ΔEX3^). (**D**) TCF/LEF reporter activity using *CTNNB1* siRNA in HEC-1-A cells. (**E**) Immunoblot validation of siRNA knockdown of β-catenin in HEC-1-A cells. TCF/LEF reporter activity with or without *Xenopus* β-catenin^ΔEX3^ transfection in **(F)** *NT5E* WT HEC-1-A cells and Ishikawa cells or (**G**) *NT5E* WT and *NT5E* KO HEC-1-A cells. (**H**) Immunofluorescence showing CD73 is localized to the membrane in Ishikawa cells overexpressing AdV- transduced *NT5E*. Membrane localization of CD73 in HEC-1-A cells serves as a positive control. Images are cropped from originals (originals are in Supplemental Figure 2) using BZ-X800 analyzer software (Keyence). Cropping was intended to more easily highlight membrane localization of CD73. All images were cropped to the same size from a 20X image. Scale bars 20 um. (**I**) TCF/LEF reporter activity in Ishikawa cells with or without *Xenopus* β-catenin^ΔEX3^ transfection and AdV-transduced *NT5E*. (**D**, **F, G, I**) Graphs are pooled data from *n* = 3 independent experiments (**D**) or *n* = 2 independent experiments (**F, G, I**). Data represent the mean ± SEM. *P < 0.05, ****P < 0.0001; 2-way ANOVA with Sidak’s post test.

### CD73 restrains exon 3 *Xenopus* mutant β-catenin transcriptional activity

With the successful validation of the TCF/LEF reporter assay in EC cells, we next examined whether CD73 loss alters the transcriptional activity of mutant β-catenin. We measured *Xenopus* β-catenin^ΔEX3^-responsive reporter gene activity in *NT5E* wild-type HEC-1-A cells (*NT5E* WT) and an *NT5E*-/- HEC-1-A cell line (*NT5E* KO) generated by CRISPR/Cas9-mediated editing of the *NT5E* gene. Ectopically expressed *Xenopus* β-catenin^ΔEX3^ induced ∼10-fold higher levels of TCF/LEF reporter activity in *NT5E* KO cells when compared with *NT5E* WT cells (Figure 2G), indicating that CD73 limits transcriptional activity of exon 3 mutant β-catenin. CD73 status and equivalent expression of *Xenopus* β-catenin^ΔEX3^ in *NT5E* WT and *NT5E* KO cells was confirmed by immunoblotting (Figure 2G).

Next, we tested the effect of ectopically expressed CD73 on β-catenin-driven TCF/LEF activity in Ishikawa cells (which lack endogenous CD73). We used an *NT5E* adenoviral vector (AdV) to reconstitute CD73 expression in Ishikawa cells. We ensured that Ishikawa cells were reconstituted with a level of CD73 expression that was equivalent to endogenous CD73 levels in HEC-1-A *NT5E* WT cells (Supplemental Figure 2A). As shown in (Figure 2H), Ishikawa cells transduced with *NT5E* AdV showed a pattern of membrane-localized CD73, which was similar to the distribution of endogenous CD73 in HEC-1-A *NT5E* WT cells (Supplemental Figure 2C).

We measured the effect of reconstituted CD73 on *Xenopus* β-catenin^ΔEX3^-dependent transcriptional activity of the TCF/LEF reporter construct. Consistent with a role of CD73 in restraining mutant β-catenin activity, *Xenopus* β-catenin^ΔEX3^-driven TCF/LEF reporter activity was reduced 2-fold in Ishikawa cells reconstituted with CD73 when compared with control cultures (Figure 2I). Reconstitution of CD73 protein levels, and equivalent expression of *Xenopus* β-catenin^ΔEX3^ expression in control and CD73-complemented cells was confirmed by immunoblotting (Figure 2I, Supplemental Figure 2B). Taken together, these data suggest a critical role for CD73 in controlling mutant β-catenin transcriptional activity in EC.

### Selection of patient-specific β-catenin mutants from public databases for study in EC cells

Having demonstrated that CD73 can restrain the transcriptional activity of *Xenopus* β- catenin^ΔEX3^, a 4-residue mutant, we asked whether CD73 restrains the activity of exon 3 *CTNNB1* mutants that are found in human EC with only one mutated codon. An additional consideration was the wide variety of missense mutations in exon 3 of *CTNNB1* reported for EC^16,17,40,41^. To select the most patient-relevant mutants to develop expression constructs, mutational data was pooled from five patient cohorts, and variety and frequency of the mutations graphed (Figure 3A). As expected, the most commonly mutated codons were S37 and S33, which are residues that when phosphorylated by glycogen synthase kinase-3β initiate the degradation of β-catenin^39,42–45^. Non-serine/threonine residues D32 and G34 were also highly mutated. Mutations at these sites interfere with β-catenin ubiquitination and subsequent degradation by reducing both β-catenin binding to E3 ligase β-TrCP as well as β-TrCP- mediated ubiquitination of β-catenin^46,47^. We selected 7 mutants to test for negative regulation by CD73: phosphorylation site mutants S33F, S33Y, S37C, S37F, and S45F, and non-phosphorylated site mutants D32N and G34R. We additionally included WT *CTNNB1*, as overexpression of β-catenin is oncogenic and is found in EC^16,48–50^.

**Figure 3.**
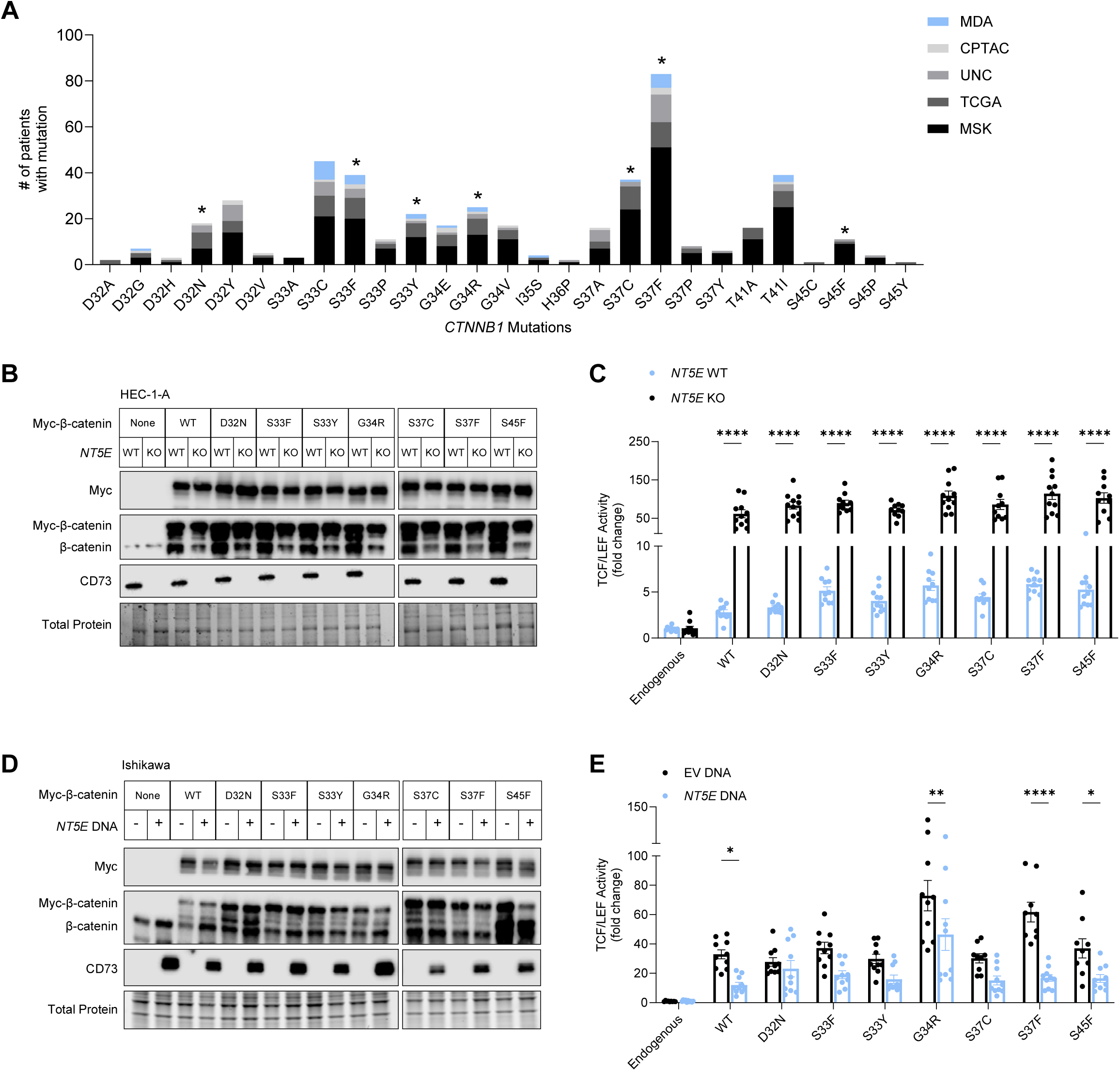
CD73 restrains transcriptional activity of patient-specific β-catenin mutants. (**A**) Variety and frequency of missense mutations in exon 3 of *CTNNB1* in endometrial cancer patients, collected from 5 patient cohorts (The Cancer Genome Atlas (TCGA), Memorial Sloan Kettering Cancer Center (MSK), Clinical Proteomic Tumor Analysis Consortium (CPTAC), the University of North Carolina (UNC), and the University of Texas MD Anderson Cancer Center (MDA). Asterisks indicate patient-specific *CTNNB1* mutations which expression vectors were developed and used in our studies. **(B, D)** Immunoblots of myc-tagged wildtype (WT) and patient-specific β-catenin mutants and CD73 expression in HEC-1-A and Ishikawa cells. **(C, E)** TCF/LEF reporter activity of WT and patient-specific β-catenin mutants in HEC-1-A and Ishikawa cells. (**E**) Ishikawa cells were transfected with AdV *NT5E* or AdV empty vector DNA. (**B**) and (**D**) are representative immunoblots from two independent experiments. (**C**) and (**E**) are pooled data from *n* = 2 independent experiments of 5-6 replicates per experiment. Data represent the mean ± SEM. *P < 0.05, ** P < 0.01, ***P < 0.0005, ****P < 0.0001; 2-way ANOVA with Sidak’s post test.

### CD73 restrains the transcriptional activity of patient-specific exon 3 β-catenin mutants

Using plasmid transfection, we successfully expressed C-terminal MYC-tagged forms of all 7 patient-specific β-catenin mutants (and wildtype (WT) β-catenin) in HEC-1-A *NT5E* WT and *NT5E* KO cells (Figure 3B). Similar to our data with *Xenopus* β-catenin^ΔEX3^, CD73 loss led to increased TCF/LEF reporter activity ∼20-fold in response to WT β-catenin and all seven patient-specific β-catenin mutants (Figure 3C). Thus, CD73 downregulation not only increases mutant β-catenin activity in *CTNNB1* mutant tumors but potentially promotes the oncogenic activity of WT β-catenin in EC.

In a reciprocal experiment, ectopic expression of CD73 in Ishikawa cells (which lack endogenous CD73; Figure 3D) led to the repression of β-catenin-driven TCF/LEF activity by 2-4-fold with patient relevant mutants, specifically G34R, S37F, and S45F, as well as WT β-catenin (Figure 3E). CD73 to restrain WT β-catenin transcriptional activity is consistent with our previous studies showing that CD73 can control the localization of non-exon 3 β-catenin to the cell membrane^31^. Thus, CD73 likely limits the oncogenic activity of both aberrantly expressed WT β-catenin and exon 3 β-catenin in EC. The reduction of β-catenin activity by ectopic expression of CD73 in some mutants in the Ishikawa cells is evidence of β-catenin mutant-specific differences in EC.

### CD73 sequesters exon 3 mutant β-catenin to the cell membrane

We previously reported that CD73-generated adenosine promotes epithelial integrity by moving cytoplasmic E-cadherin and wildtype β-catenin to the cell membrane^31^. Given that exon 3 encodes a region on the N-terminus of β-catenin, which is separate from the region where β-catenin binds with E-cadherin^33–36^, we hypothesized that CD73 may restrain mutant β-catenin transcriptional activity by sequestering it to the cell membrane.

As proof of principle, we first assessed the nuclear localization of *Xenopus* β-catenin^ΔEX3^ in HEC-1-A cells treated with *NT5E*-directed siRNA (or control non-targeting siRNA). Consistent with our hypothesis, knockdown of *NT5E* led to increased nuclear *Xenopus* β-catenin^ΔEX3^ (Figure 4A-4C) as detected by immunofluorescence microscopy. To further test our hypothesis, cellular fractionations and immunoblotting were performed to compare the levels of mutant β-catenin in the different cellular compartments of *NT5E* WT and KO HEC-1-A cells. Fully consistent with the results of our immunofluorescence microscopy experiments, we observed that in *NT5E* KO HEC-1-A cells, mutant β-catenin membrane levels were decreased up to 2-fold, whereas nuclear and chromatin-bound levels were increased compared with *NT5E* WT HEC-1-A cells (Figure 4D-4F, Supplemental Figure 4A-4C).

**Figure 4.**
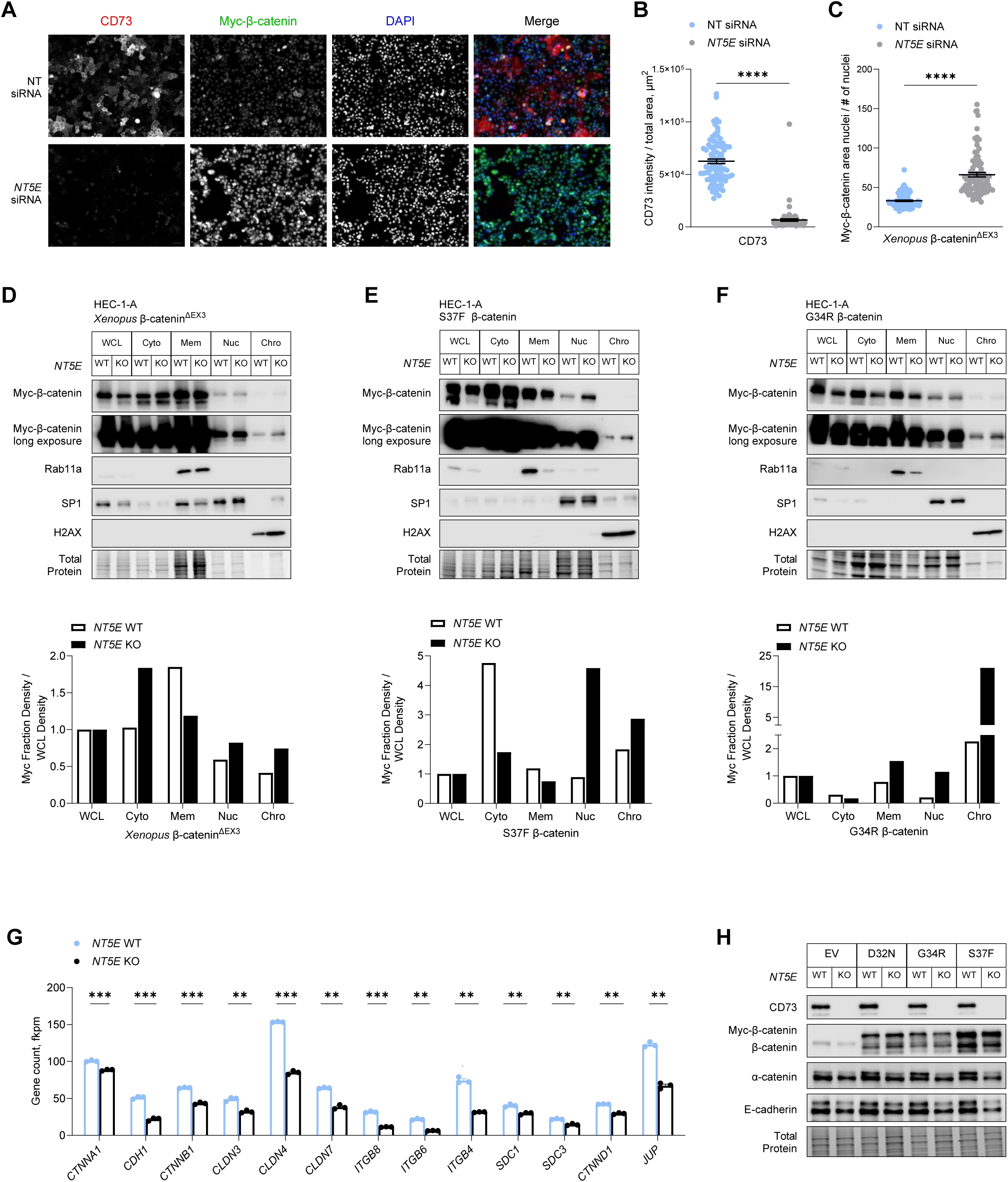
CD73 sequesters exon 3 mutant β-catenin to the cell membrane. (**A-C**) HEC-1-A cells were transfected with *Xenopus* β-catenin^ΔEX3^ and NT (non-targeting) or *NT5E* siRNA and cultured in 1% O_2_ and 5% CO_2_ for 48 hours. **(A)** Representative immunofluorescence images used in quantifying nuclear localization of myc-β-catenin^ΔEX3^. Scale bar: 50 µm. Fluorescence intensity was determined with BZ-X800 Analyzer Macro cell count software (Keyence). (**B)** Validation of CD73 knockdown and (**C**) nuclear fluorescence intensity of myc-β-catenin^ΔEX3^. (**D-F**) Representative immunoblots of *n* = 2 independent experiments of cellular fractionations from *NT5E* WT and *NT5E* KO HEC-1-A cells. Cells were transfected with (**D**) *Xenopus* β-catenin^ΔEX3^ or patient-specific β-catenin mutants **(E)** S37F or **(F)** G34R. Graphs show densitometry for myc-β-catenin mutant expression for each cellular fraction normalized to myc-β-catenin mutant expression in the whole cell lysate (WCL). Cellular fraction markers: Rab11a (membrane), SP1 (nuclear), and H2AX (chromatin). (**G**) mRNA and (**H**) protein expression of differentially expressed cell-cell adhesion components in *NT5E* WT and *NT5E* KO HEC-1-A cells. Data represent the mean ± SEM. ****P < 0.0001, Mann-Whitney test; **P < 0.01, ***P < 0.005; Welch t-test.

A 2-fold increase in nuclear mutant β-catenin and a 3-fold increase in chromatin-bound mutant β-catenin were seen with *Xenopus* β-catenin^ΔEX3^ (Figure 4D, Supplemental Figure 4A) and patient-specific β-catenin mutants, S37F (Figure 4E, Supplemental Figure 4B) and G34R (Figure 4F, Supplemental Fig 4C). We accounted for the random variability in mutant β-catenin expression in *NT5E* KO vs. WT cells (Figure 4D-4F and Supplemental Figure 4A-4C) in our calculations by normalizing the expression of mutant β-catenin in each cell compartment to mutant β-catenin expression in the whole cell lysate (WCL) for each cell type. Together, these data suggest that CD73 restrains mutant β-catenin activity by sequestering it to the cell membrane.

Consistent with this interpretation, we show in Supplemental Figure 3 that patient-specific mutant β-catenin does bind with E-cadherin in *NT5E* KO and WT HEC-1-A cells. Here we observed that total E-cadherin levels in *NT5E* KO cells were significantly lower than *NT5E* WT cells (Figure 4H and Supplemental Figure 3). The difference in E-cadherin levels between *NT5E* KO vs. WT HEC-1-A cells prompted us to investigate the expression of other cell-cell adhesion genes. RNA-seq data revealed that *NT5E* KO cells compared to *NT5E* WT cells were deficient in the expression of many genes essential to the assembly and stabilization of cell-cell adhesions (Figure 4G), including α-catenin (Figure 4G and 4H), which stabilizes E-cadherin-β-catenin complexes at the cell membrane by binding the actin filaments^51–53^. These data are in accordance with our previous work showing that CD73 is essential for epithelial integrity^31^. Therefore, the ability of CD73 to maintain epithelial cell-cell adhesions is likely one mechanism by which CD73 restrains mutant β-catenin oncogenic activity.

### CD73 restrains mutant β-catenin transcriptional activity through adenosine receptor A1 activity

To test whether CD73 restrains β-catenin transcriptional oncogenic activity by adenosine receptor signaling, we generated HEC-1-A cells with CRISPR-Cas9 deletion of adenosine receptors, A_1_R (*ADORA1*) and A_2B_R (*ADORA2B*) (Figure 5A-B). HEC-1-A cells largely express two of the four adenosine receptors (Figure 5A). Notably, unlike *NT5E* KO HEC-1-A cells, which cell-cell adhesions are globally dysregulated (e.g., E-cadherin and other cell-cell adhesion gene expression are downregulated compared with *NT5E* WT; Figure 4G), *ADORA1* KO and *ADORA2B* KO HEC-1-A cells expressed similar E-cadherin levels when compared to WT HEC-1-A cells (Figure 5C, Supplemental Figure 5A). Thus, *ADORA1* and *ADORA2B* KO cells, retain intact cell-cell adhesions.

**Figure 5.**
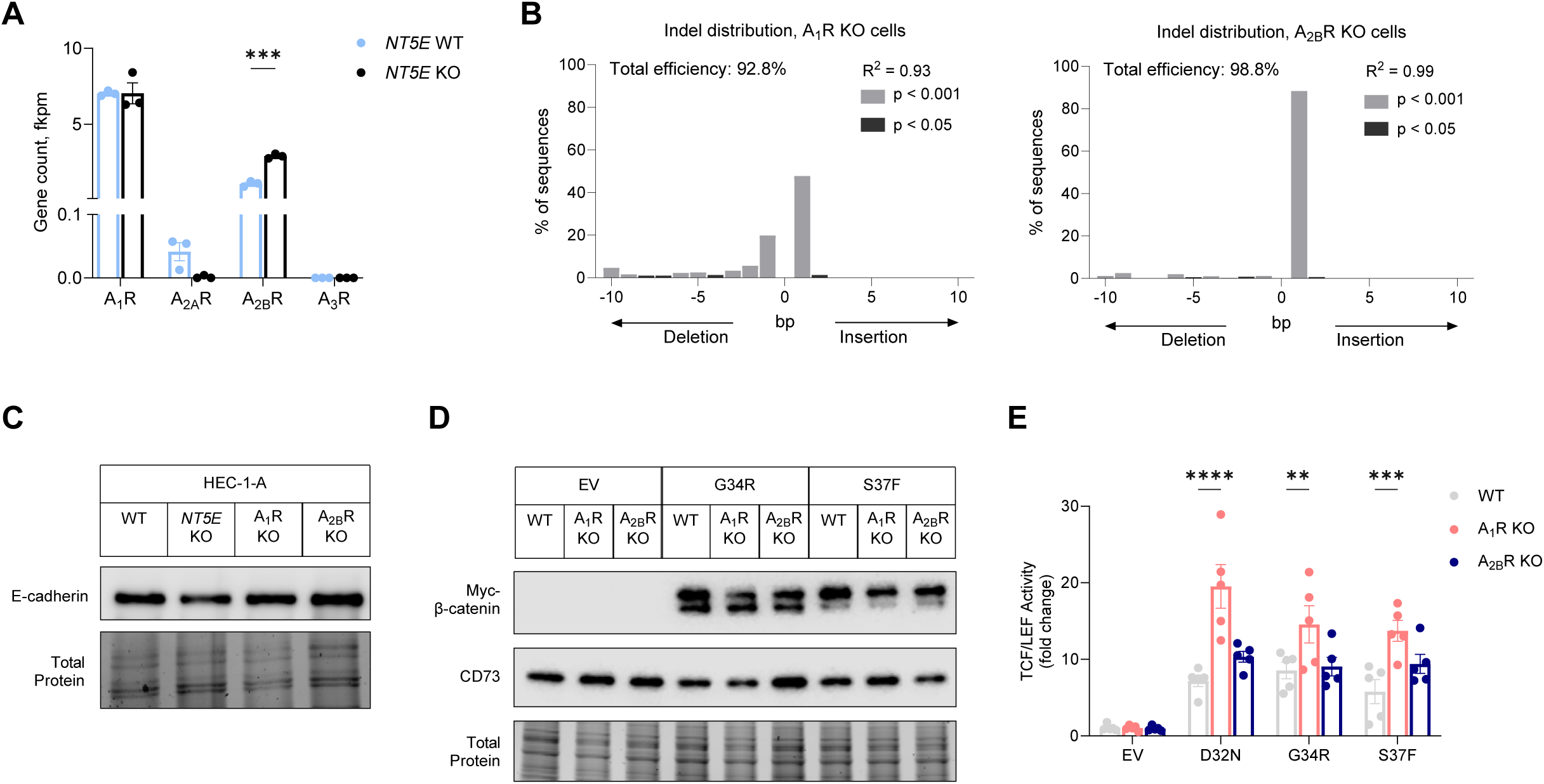
Adenosine A1 receptor signaling restrains the transcriptional activity of patient-specific β-catenin mutants. (**A**) mRNA expression of adenosine receptors in *NT5E* WT and *NT5E* KO HEC-1-A cells. (**B**) TIDE analysis for *ADORA1* KO and *ADORA2B* KO CRISPR-Cas9 deleted HEC-1-A cells. (**C**) Immunoblots showing epithelial integrity (E-cadherin as a readout) is not disrupted in *ADORA1* KO and *ADORA2B* KO HEC-1-A cells. (**D**) Approximate equivalent expression of myc-G34R β-catenin and myc-S37F β-catenin in WT, *ADORA1* KO, and *ADORA2B* KO HEC-1-A cells. (**E**) TCF/LEF reporter activity in cells transfected with empty vector (EV), D32N, G34R, or S37F β-catenin. Each dot represents one technical replicate. Graph is representative of *n* = 3 independent experiments; other independent experiments are shown in Figure S6. (**A-D**) A_1_R = *ADORA1*, A_2A_R = *ADORA2A*, A_2B_R = *ADORA2B*; A_3_R = *ADORA3*. Data represent mean ± SEM. *P < 0.05, **P < 0.01, ***P < 0.0005; 2-way ANOVA with Dunnett’s post test.

We measured TCF/LEF reporter activity in response to ectopically expressed patient-specific β-catenin mutants (D32N, G34R, and S37F) in *ADORA1* KO, *ADORA2B* KO, and WT HEC-1-A cells. The mutants were evenly expressed in the different cell lines (Figure 5D). D32N β-catenin led to a ∼7-fold increase in TCF/LEF reporter gene activity relative to empty vector in WT cells. However, in *ADORA1* KO cells, D32N β-catenin induced a ∼20-fold increase in TCF/LEF reporter activity. In *ADORA2* KO cells, the magnitude of mutant β-catenin-driven TCF/LEF activity was similar to WT HEC-1-A cells. Qualitatively similar results were observed for the G34R and S37F β-catenin mutants (Figure 5E and Supplemental Figure 5B-C). Together these data show that similar to *NT5E*-deficiency, *ADORA1* loss leads to de-repression of mutant β-catenin transcriptional activity.

Previously, we showed CD73-A_1_R signaling limits disease aggressiveness in EC by protecting cell-cell adhesions through increasing the localization of WT β-catenin to the membrane^31^. Accordingly, CD73-A_1_R signaling in exon 3 β-catenin mutant tumors likely is functioning in a similar way, and therefore, restrains β-catenin mutant transcriptional activity by redistributing mutant β-catenin to the cell membrane.

### Loss of CD73 induces pro-tumor Wnt/β-catenin target gene expression

The robust TCF/LEF transcriptional activity observed in *NT5E* KO HEC-1-A cells (seen in Figure 2E and Figure 3C) led us to perform RNA-seq. Our goal was to identify gene targets of mutant β-catenin that occur only with the loss of CD73. Patient-specific mutants, D32N, G34R, and S37F, and empty vector were transfected into *NT5E* KO and WT HEC-1-A cells. Similar expression of the mutants in *NT5E* KO vs. WT cells were validated by immunoblot and transcript levels (Supplemental Figure 6A-6B). Expression of β-catenin target genes (such as *TCF7* and *AXIN2*) confirmed that β-catenin mutants in *NT5E* KO and WT cells were transcriptionally active (Supplemental Figure 6D-6E). As expected, *NT5E* KO vs. WT samples showed the greatest separation by the principal component analysis. For *NT5E* KO samples, the β-catenin mutants showed separation from the empty vector *NT5E* KO samples (Figure 6A). *NT5E* WT samples were similar with the exception that *NT5E* WT D32N samples which clustered closely with empty vector *NT5E* WT samples (Figure 6A).

**Figure 6.**
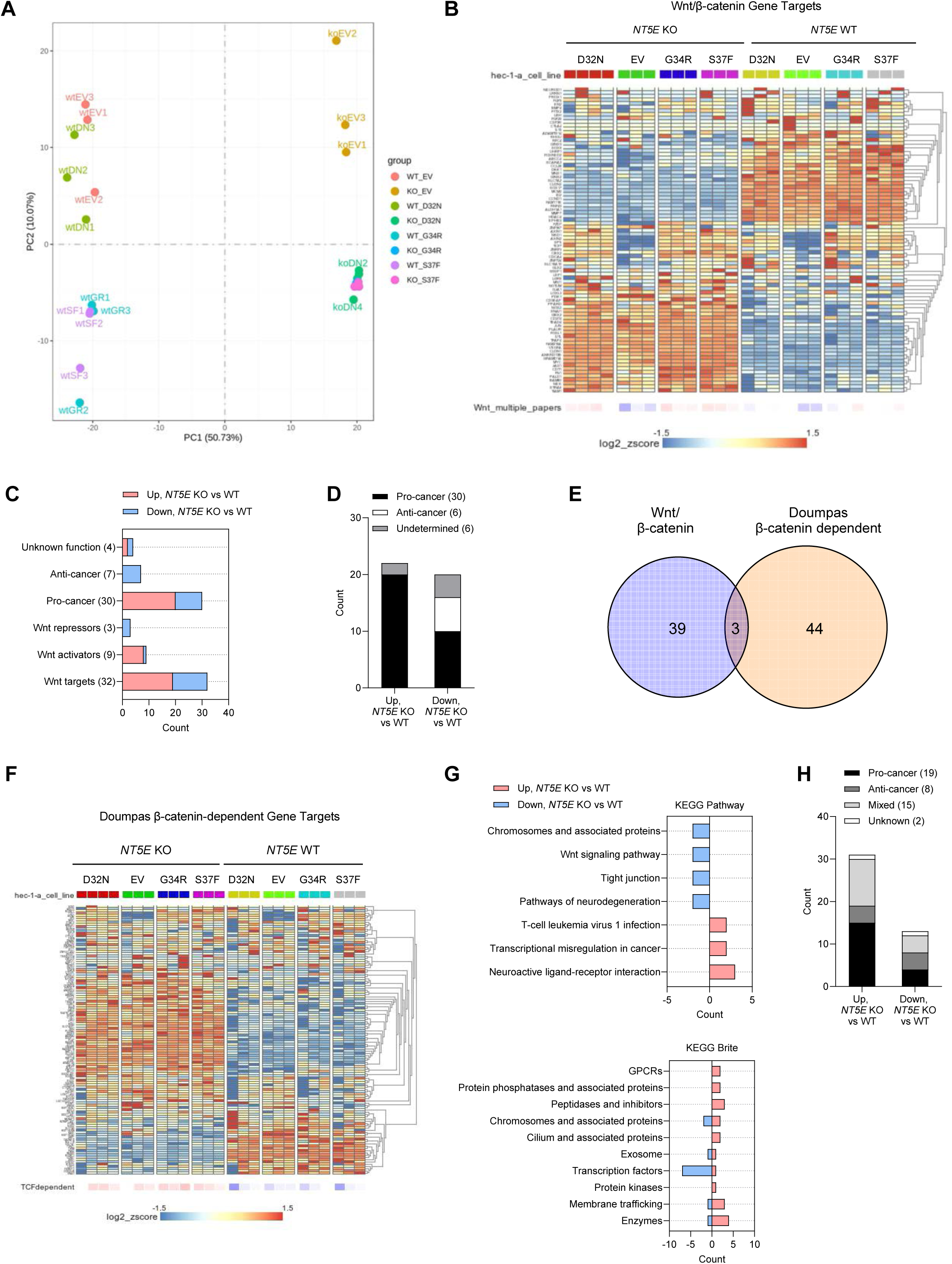
Loss of CD73 results in an oncogenic WNT/β-catenin gene signature. (**A**) PCA plot for RNA-seq data from *NT5E* WT vs. *NT5E* KO HEC-1-A cells expressing empty vector (EV) or different β-catenin mutants. (**B**-**D**) RNA-seq data analyses using literature-derived Wnt/β-catenin signaling target genes. (**B**) Heatmap showing global differences in Wnt/β-catenin signaling target genes between *NT5E* WT vs. *NT5E* KO. (**C**) Analysis of genes from (**B**) that were significantly different between *NT5E* KO EV vs. *NT5E* WT EV. Genes were placed in categories based on Wnt function and literature-reported oncogenic (pro-tumor) or tumor suppressor (anti-cancer) activity. Total number of genes in each category is indicated in parentheses. (**D**) Combined pro-cancer and anti-cancer-associated genes by upregulation or downregulation status in *NT5E* KO EV vs. *NT5E* WT EV samples. (**E**) Venn diagram of Wnt/β-catenin signaling target genes and Doumpas Wnt/β-catenin-dependent target genes, significantly disregulated between *NT5E* WT vs *NT5E* KO cells. (**F-H**) RNA-seq data analyses using Doumpas Wnt/β-catenin-dependent target genes. (**F**) Heatmap showing global differences in Doumpas Wnt/β-catenin-dependent target genes between *NT5E* WT vs. *NT5E* KO. (**G**) KEGG pathway and Brite analyses of genes significantly altered in (**F**) in *NT5E* KO EV vs. *NT5E* WT EV samples^123^. (**H**) Combined pro-cancer and anti-cancer-associated genes significantly altered in *NT5E* KO EV vs. *NT5E* WT EV samples.

Unexpectedly, we observed substantial Wnt/β-catenin target gene expression changes between *NT5E* WT vs. KO HEC-1-A samples, regardless of mutant β-catenin or empty vector expression (Figure 6B). Using a list of canonical Wnt/β-catenin pathway target genes, compiled from various publications, we found that of 76 genes assessed, 20 were significantly downregulated and 22 were significantly upregulated in *NT5E* KO cells compared to *NT5E* WT HEC-1-A cells (Supplemental Tables 3 and 4). Genes upregulated in *NT5E* KO vs. WT cells were predominately pro-cancer genes (Figure 6C and 6D). Given the strong oncogenic Wnt/β-catenin target gene signature of *NT5E* KO cells, we used a second β-catenin dependent gene list by Doumpas et. al. (*EMBO J, 2019)* which reported genes targets of β-catenin based on chromatin immunoprecipitation (ChIP) and TCF deletion in cells^54^. Only 7 genes were found to overlap between these two gene lists, 3 of which showed significant differences in *NT5E* KO vs. *NT5E* WT cells (Figure 6E). Similar to our literature-derived list (Figure 6B), of the 112 unique Doumpas genes there were substantial gene expression changes between *NT5E* WT vs. KO samples, regardless of β-catenin mutants or empty vector (Figure 6F). KEGG analyses were performed on genes that were significantly upregulated (*n* = 31) and downregulated (*n* = 13) (Supplemental Tables 5 and 6). Upregulated genes were involved in pathways such as transcriptional misregulation in cancer, neuroactive ligand-receptor interactions, and T-cell leukemia virus 1 infection (Figure 6G). Of the upregulated genes, half were associated with pro-cancer activity (Figure 6H), and a similar number showed mixed activity (Figure 6H). We selected three of these genes (*SOX17*, *JUN,* and *FOSL1*) to validate by protein expression (Supplemental Figure 7). Together, these data reveal that CD73 loss alone is capable of driving pro-tumor Wnt/β-catenin target gene expression regardless of mutant β-catenin expression.

### Zinc finger transcription factors and non-coding RNAs are gene targets for patient-specific β-catenin mutants in *NT5E* KO cells

Despite strong differences in gene expression between *NT5E* KO vs. WT HEC-1-A cells, several differentially expressed genes were identified with the different patient-specific mutants that occurred only with CD73 loss: D32N *NT5E* KO cells (*n* = 143), G34R *NT5E* KO cells (*n* = 78), and S37F *NT5E* KO cells (*n* = 80) (Figure 7A). The majority of the protein coding genes were non-Wnt/β-catenin target genes and were largely downregulated (Figure 7B, 7E, and Supplemental Table 7). KEGG analyses revealed that all mutants commonly showed downregulation of zinc finger transcription factors, such as *ZFN708*, *ZNF782*, and *ZNF112* (Figure 7D; herpes simplex virus 1 infection and transcription factors; Figure 7E). Various studies show that ZNFs can exhibit anti-tumor behavior by downregulating Wnt signaling and cell growth, promoting epithelial-to-mesenchymal transition (EMT), ribosome biogenesis, and NF-κB signaling, as well as inducing apoptosis^55–57^.

**Figure 7.**
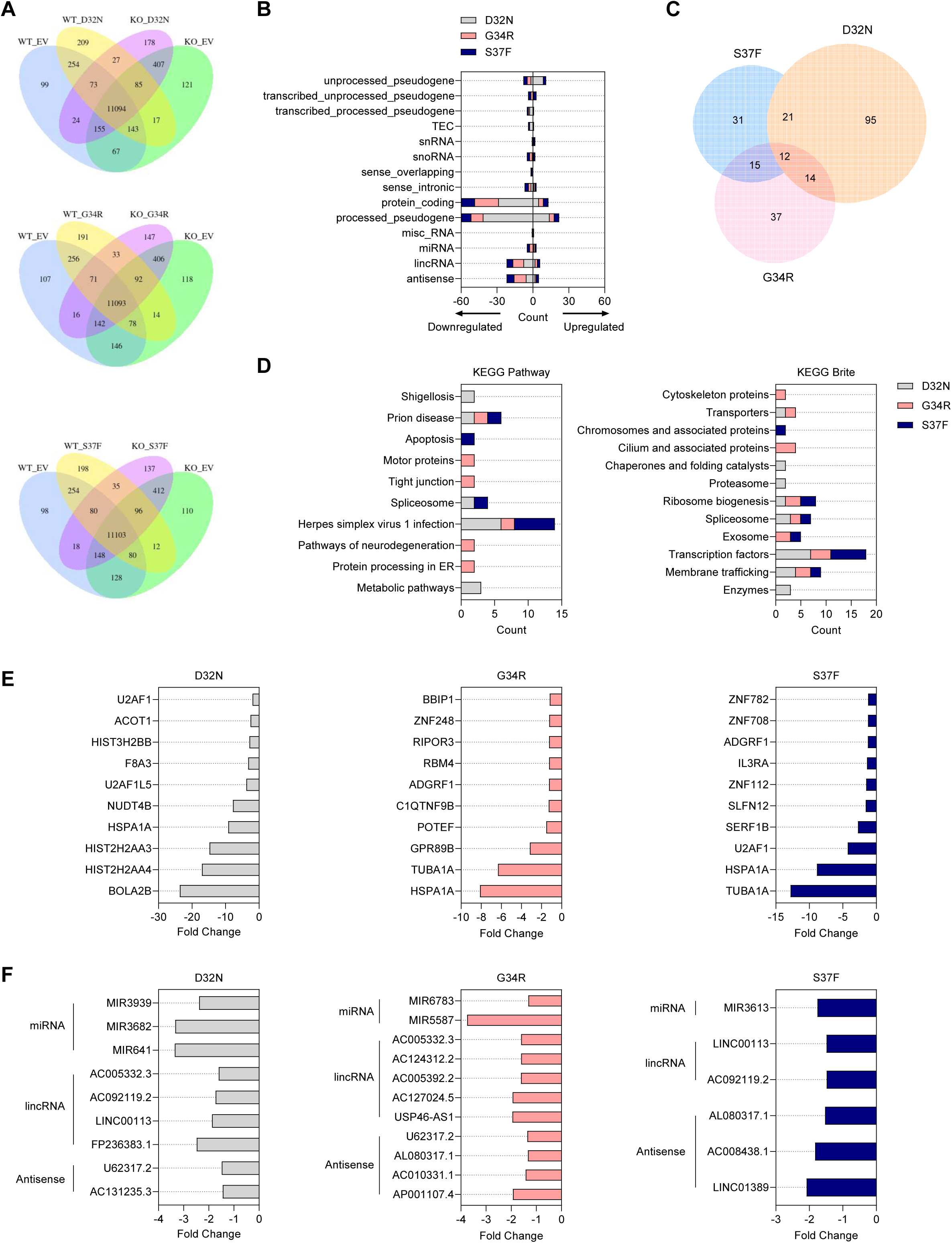
Mutant-specific transcriptional expression in *NT5E* KO HEC-1-A cells. (**A-F**) RNA-seq data from *NT5E* WT vs. *NT5E* KO HEC-1-A cells expressing empty vector (EV) or different β-catenin mutants. (**A**) Venn diagrams of differentially expressed genes between one β-catenin mutant vs. EV in *NT5E* WT and *NT5E* KO samples. (**B**) Classification of 301 altered genes from each mutant in *NT5E* KO cells. These are represented in a (**A**) as *n* = 143 genes for D32N; *n* = 78 genes for G34R; *n* = 80 genes for S37F. (**C**) Venn diagram of 301 selected genes showing overlap between genes targeted by various mutants. Venn diagram generated with Meta-Chart Venn Diagram Maker Online webtool, then re-made with calculated proportions in GraphPad Prism. (**D**) KEGG Pathway and Brite analyses for protein-coding mutant-specific genes differentially expressed in *NT5E* KO cells^123^. (**E**) Protein-coding mutant-specific genes differentially expressed in *NT5E* KO cells; for each mutant, 10 genes with largest fold change difference in (mutant) *NT5E* KO vs EV *NT5E* KO are shown. (**F**) miRNA, lincRNA, and antisense RNA most differentially expressed by each mutant in *NT5E* KO cells (*n* = 26 genes; extra information in Supplemental Table 7).

Consistent with the consideration that different patient-specific β-catenin mutants likely induce different gene expression programs which results in different biology in EC, we found individual mutants showed gene expression differences preferentially for metabolic pathways/enzymes (D32N), neurodegeneration pathways/cytoskeleton and cilium associated proteins (G34R), or apoptosis (S37F) (Figure 7D). Interestingly, processed pseudogenes and non-coding RNAs (ncRNAs), including miRNAs, lincRNAs, and antisense RNAs, were also identified to be differentially expressed, largely downregulated, and mutant-specific with the different patient-specific mutants with CD73 loss (Figure 7B, 7F). Various studies have reported that ncRNAs can have various tumor-promoting or tumor-suppressing roles in cancer, and much more information needs to be elucidated to fully understand these roles^58–60^. However, several of our identified downregulated ncRNAs were found to have anti-cancer and or “mixed” functions, including *MIR3613*, *AC010331.1,* and *MIR641* (Supplemental Table 7). Thus, mutant β-catenin downregulating anti-tumor ncRNAs could be a potential mechanism for promoting the aggressiveness of mutant β-catenin EEC. Altogether these data show β-catenin mutant tumors gain unique, non-Wnt gene expression changes with CD73 loss, and they provide evidence for mutant-specific differences in EC.

## Discussion

We have identified CD73 as a novel regulator of a major oncogene in EC. Using human tumors and genetic approaches, we showed that CD73 restrains the oncogenic transcriptional activity of exon 3 β-catenin mutants in EC. Mechanistically, we provided evidence that CD73 limits mutant β-catenin nuclear and chromatin localization by sequestering mutant β-catenin to the cell membrane, likely through CD73-A_1_R signaling. Additionally, we revealed that CD73 loss associates with recurrence in patients with exon 3 *CTNNB1* mutant EC, and that CD73 loss alone in EC cells promotes pro-tumor, Wnt/β-catenin target gene expression.

Recent studies demonstrate the value of molecular testing when guiding clinical decisions for patients with EC^26,61–69^. Four molecular subtypes for EC have been identified^26,70,71^, and tumors with *CTNNB1* mutation largely fall into the no specific molecular profile (NSMP) subtype. NSMP tumors are clinically challenging as they are a mix of indolent and high-risk disease with no clear prognostic markers. *CTNNB1* has been unsuccessful as a reliable marker despite high expectations. Our work strongly suggests that CD73 may improve the power of *CTNNB1* mutation as a predictive biomarker of recurrence. When tumors were individually separated by *NT5E* expression, we showed the lower quartile cutoff to capture 86% of patients within this group that recurred. Large scale studies will be necessary to determine the clinical value of this finding. While we focused on tumors with exon 3 *CTNNB1* mutations, investigating CD73 in NSMP tumors with overexpression or activated Wnt signaling may also prove valuable, as we showed that CD73 can regulate the transcriptional activity of both WT and exon 3 β-catenin mutants. Studies like these will be important given recent data showing that adjuvant therapy can improve outcomes for patients with tumors showing aberrant β-catenin expression or harboring *CTNNB1* mutations^72,73^. Currently, clinical surveillance is the standard of care for these patients.

Our discovery that CD73 localizes mutant β-catenin to the cell membrane provides a novel regulatory mechanism for a major oncogene. Despite advances in small molecule design, natural product isolation, and miRNA utilization, anti-β-catenin therapies have not translated into the clinic^74^. Thus, efforts to uncover modulators of β-catenin activity are vital for identifying novel vulnerabilities that can be exploited for improving patient outcomes. We anticipate the regulatory action by CD73 is cancer-type specific, as CD73 is downregulated in EC^31,32^, whereas in other human cancers, CD73 is overexpressed^75–78^. Exon 3 *CTNNB1* mutations are also found in hepatocellular and prostate carcinomas^79–81^. Both tumors are reported to downregulate CD73 expression or its enzymatic activity^82–84^. Therefore, an oncogenic mechanism involving de-repression of mutant β-catenin by CD73 loss may be particularly relevant to these tumors.

With the frequency of hotspot exon 3 *CTNNB1* mutations in EC, the oncogenic mechanism of mutant β-catenin is often attributed to the Wnt/β-catenin signaling axis. Less attention has been given to the cell-cell adhesion function of β-catenin, despite decades of studies demonstrating that the two functions are independent and that mutations in exon 3 of *CTNNB1* do not impact β-catenin binding with E-cadherin and reaching the cell membrane^2,85–87^. Our work is consistent with these studies and importantly demonstrates that cell-cell adhesions in EC cells are molecular sinks for oncogenic β-catenin. The critical nature of cell-cell adhesions (specifically E-cadherin binding) to control the oncogenic activity of β-catenin is underscored by studies showing that binding of β-catenin to E-cadherin prevents β-catenin nuclear localization and β-catenin/LEF-1-mediated transactivation^2,85,86^, which β-catenin binding with LEF-1 can be out-competed by E-cadherin^2^, as well as studies showing that E-cadherin can limit the transforming properties of activating β-catenin mutations^2,87,88^.

Whether the ability of CD73 to sequester mutant β-catenin to the membrane is entirely dependent on E-cadherin expression is unclear. E-cadherin expression in low grade, early stage EC is variable; E-cadherin staining ranges from 5-95% for these tumors^89^. The status of E-cadherin in the human tumors used in our study was not assessed. E-cadherin is part of a large family of cadherin proteins, whereby N-cadherin is structurally similar to E-cadherin and is highly expressed in low grade, early stage EEC, including tumors with *CTNNB1* mutations^16^. While N-cadherin is largely known for promoting cancer cell migration, N-cadherin also promotes the formation of cell-cell adhesions. Similar to E-cadherin, N-cadherin binds with β-catenin and can block β-catenin/LEF-1- mediated transactivation^90,91^. N-cadherin ligation and the initiation of F-actin branching for cell movement is inhibited when forming cell-cell contacts^92^. Additionally, post-translational modifications and the presence of binding factors (such as Fibroblast Growth Factor Receptor (FGFR)) influence whether N-cadherin promotes migration or stabilizes cell-cell adhesions^92–94^. Thus, in some tumors, N-cadherin may serve as a molecular sink for mutant β-catenin, and therefore E-cadherin expression may not be a reliable readout of CD73 to restrain mutant β-catenin. Notably, tumor differentiation significantly affects E-cadherin expression in EC, but has no impact on N-cadherin levels^95^.

While cell-cell adhesions are advantageous for suppressing the activity of oncogenes, these structures can become saturated. Saturation of adherens junctions by oncogenic β-catenin binding to E-cadherin has been demonstrated in studies using murine models of intestinal cancer^87^. Additionally, recent evidence suggests that fold change – not absolute levels – of β-catenin can dictate Wnt signaling^96^. Accordingly, in addition to the expression level of E-cadherin or other cadherins (e.g., N-cadherin), the expression level of mutant β-catenin in EC likely also determines the amount of mutant β-catenin that can be sequestered at the cell membrane. These factors may explain why some patients with *CTNNB1* mutant tumors that retained CD73 expression had disease recurrence (Figure 1).

The impact of CD73 loss is likely more than losing the localization of mutant β-catenin to the membrane. For example, CD73 deletion in HEC-1-A cells resulted in the downregulation of *SOX17* and upregulation of *JUN* and *FOSL1* expression. *SOX17* is a negative regulator of β-catenin/TCF transcription and inhibits EC progression by inactivating Wnt/β-catenin-mediated proliferation and EMT^97,98^. Similar to TCF, *JUN* and *FOSL1* are Wnt target transcription factors that complex with β-catenin and drive target gene expression and tumor progression^99–103^. These transcriptomic changes likely help explain our unexpected observation that CD73 loss alone, regardless of empty vector or β-catenin mutant expression, in HEC-1-A cells led to a strong induction of pro-tumor Wnt/β-catenin target gene expression. Notably, endogenous β-catenin levels in *NT5E* KO HEC-1-A cells are downregulated. Thus, while these data are less clear, they suggest that the amount of nuclear β-catenin in EC may not be as important as the robustness of the nuclear transcriptional activity of mutant β-catenin and the selection of the genes that are targeted. Here the difference would be caused by the absence or presence of negative regulators that would compete with β-catenin-TCF/LEF transcription sites and the added presence of multiple binding proteins (e.g., transcription factors such as *JUN* and *FOSL1*) to drive gene expression^98,104^. Overall, it is interesting to consider that the downregulation of CD73 may be an additional mechanism by which endometrial tumors to gain oncogenic Wnt/β-catenin target gene expression without β-catenin mutation or overexpression.

We made a special effort to investigate different patient-specific exon 3 β-catenin mutants found in EC. Few studies have investigated different β-catenin mutants in EC biology^105,106^, and it remains largely unclear mechanistically whether different mutations may have different biology which may further explain the variability in disease aggressiveness. We observed that CD73 ectopic expression repressed TCF/LEF activity for some β-catenin mutants (G34R, S37F, and S45F) and WT β-catenin in Ishikawa cells. Unlike *NT5E* KO HEC-1-A cells that have globally dysregulated cell-cell adhesions as a consequence of CD73 loss, Ishikawa cells retain functional cell-cell adhesions, which provides an opportunity to observe mutant-specific regulation by CD73. Why certain mutants are more affected by CD73 is not clear. However, it is not unexpected given that different exon 3 *CTNNB1* mutants are demonstrated to have functional differences (e.g., proliferation and migration capacity) in other model systems and that protein structure of β-catenin mutants (e.g., electrostatic charge, polar interactions, and stability) are differentially affected by certain amino acid changes^106–109^. Notably, our studies using adenosine receptor-deficient cells, which retain E-cadherin expression similar to WT HEC-1-A levels, demonstrated that CD73-A_1_R signaling to restrain the transcriptional activity of mutant β-catenin is not merely a consequence of the low abundance of cell-cell adhesions.

We performed RNA-seq with 3 patient-specific mutants to further examine whether regulation of β-catenin transcriptional activity by CD73 could be mutant-specific. With CD73 loss, we found that the different patient-specific mutants did not further upregulate Wnt target genes. Rather, ZNF transcription factors and ncRNAs were the most frequently dysregulated genes, and these genes were downregulated. ZNFs can have anti-tumor activity, including repressing Wnt signaling and cell growth, as well as promoting EMT, ribosome biogenesis, and NF-κB signaling, and apoptosis^55–57^. Several ncRNAs that were downregulated with β-catenin mutants in *NT5E* deficient cells have anti-cancer or ‘mixed’ functions, including *MIR3613*, *AC010331.1*, and *MIR641*^110–112^. Different exon 3 β-catenin mutants showed gene expression changes that were specific to different biological processes, such as metabolic pathways/enzymes (D32N β-catenin), neurodegeneration pathways/cytoskeleton and cilium associated proteins (G34R β-catenin), or apoptosis (S37F β-catenin). Future studies will be necessary to interrogate ZNFs and ncRNAs in the biology of *CTNNB1* mutant tumors and different patient-specific β-catenin mutants.

In summary, our study identified CD73 as a novel molecular determinant of β-catenin oncogenic activity in EC and provides the first mechanistic insight that helps explain the variability in patient outcomes in exon 3 *CTNNB1* mutant EC. Detailed studies interrogating the biology of different patient-specific β-catenin mutants will be important for personalized medicine efforts.

## STAR Methods

Experimental model and study participant details

### Human Tissues

Use of human tissues was approved (LAB01-718) by the Institutional Review Board of the University of Texas MD Anderson Cancer Center (MDACC).

### Cell lines

Human endometrial cancer (HEC)-1-A cells (American Type Culture Collection, ATCC) and HEC-1-A cells with CRISPR/Cas 9 deletion of *NT5E* (*NT5E* KO) were maintained in McCoy’s 5A (Iwakata & Grace Modification) Medium (Corning) supplemented with 10% (v/v) fetal bovine serum (FBS) (Genesee Scientific) and 100 U/ml penicillin, 100 mg/ml streptomycin. Ishikawa cells were provided by Changping Zou (formerly University of Arizona) and maintained in Minimum Essential Medium (Earle’s) supplemented with 10% (v/v) fetal bovine serum (FBS) (Genesee Scientific), 100 U/ml penicillin, 100 mg/ml streptomycin (Genesee Scientific), 1 mM sodium pyruvate (Sigma Aldrich), and 0.1 mM non-essential amino acids (Lonza). Cell lines were authenticated by the Characterized Cell Line Core Facility at the University of Texas MD Anderson Cancer Center and cultured in a humidified 5% CO_2_ atmosphere at 37°C. All cell lines used are female, as endometrial cancer only affects females.

## Method Details

### Generation of A_1_R KO and A_2B_R KO cell lines

CRISPR-Cas9 plasmids for knockout of *ADORA1* or *ADORA2B* were purchased from Vector Builder (VB240110). The plasmids contained a single guide RNA targeting *ADORA1* (sgRNA sequence 5’ –TCTCCTTCGTGGTGGGACTGA– 3’) or *ADORA2B (sgRNA sequence* 5’ –CACAGGACGCGCTGTACGTGG– 3’), the sequence for *S. pyogenes* Cas9, and ampicillin and puromycin resistance. Guides were designed using the CHOPCHOP web tool^113–115^. HEC-1-A cells were transfected with 2 ug CRISPR- Cas9 plasmid for *ADORA1* or *ADORA2B*. After 48 hours, cells were selected using puromycin (Sigma) for 3 days (2 ug/ml, Day 1; 5 ug/ml, Day 2; 2 ug/ml, Day 3). After selection, cells were cultured in McCoy’s 5A media with 10% FBS without antibiotic. Cells were expanded and used at passages 4-13 for experiments.

### Tracking of indels by decomposition (TIDE)

gDNA extraction was performed with QIAGEN columns. Target-specific PCR products were generated and sequenced by Azenta for analysis of CRISPR-Cas9 editing in *NT5E* KO, *ADORA1* KO, and *ADORA2B* KO cell lines. The TIDE webtool (available for free courtesy of Eva Brinkman, Tao Chen and Bas van Steensel) was used to calculate the frequency and spectrum of genome alterations introduced by CRISPR-Cas9 editing in *ADORA1* and *ADORA2B* genes^116^.

### Constructs and reagents

Patient-specific exon 3 *CTNNB1* (β-catenin) mutant and wildtype *CTNNB1* constructs were developed and purchased from Vector Builder. The vector backbone (#VB220927) for β-catenin mutant and wild-type plasmids contained RSV and EF1A promoters and ampicillin and hygromycin resistance gene sequences. Each plasmid contains the full-length *CTNNB1* gene with or without a single nucleotide substitution to generate mutations D32N, S33F, S33Y, G34R, S37C, S37F. All vectors have 6 myc tags on the C-terminus of *CTNNB1*. Wildtype and patient-specific exon 3 β-catenin plasmids were expressed using non-viral approaches (Lipofectamine 3000; Invitrogen) in all experiments. M50 Super 8x TOPFlash (Addgene plasmid #12456), M51 Super 8x FOPFlash (TOPFlash mutant; Addgene plasmid #12457), and XE49 pt beta-catenin-myc (*Xenopus* β-catenin^ΔEX3;^ Addgene plasmid #16840) were a gift from Randall Moon^38^. *Xenopus* β-catenin^ΔEX3^ was provided to us by Pierre D. McCrea. LentiV_Neo was a gift from Christopher Vakoc (Addgene plasmid # 108101)^117^ and was used as an empty vector control in experiments with C-myc β-catenin mutant constructs. pACCMV was provided by Lilly Chiou and was used as an empty vector control in experiments with pACCMV *NT5E* plasmid. CMV-pRenilla-LUC was purchased from Promega (Cat # E2261). All plasmids were propagated in *E. coli* DH5α and purified on QIAGEN or ZymoPURE columns (Genesee). Small interfering RNAs (siRNAs) included Non-targeting: 5’-GAUCAUACGUGCGAUCAGATT-3’ (Sigma), *NT5E*, 1247: 5’-CGCAACAAUGGCACAAUUATT-3 (as previously described^31^), and *CTNNB1*: 5’- CUCAGAUGGUGUCUGCUAU-3’ (Sigma). A complete list of antibodies is provided in Supplemental Table 2.

### Design of adenoviral vectors

The *NT5E* open reading frame with an N-terminal HA-tag was cloned into the pACCMV vector^118^ and plasmid purified using the Qiagen EndoFree Plasmid Maxi Kit. Adenovirus purification and infections were performed as described previously^119^. In brief, the *NT5E* pACCMV vector or an empty pACCMV vector was co-transfected in 239T cells with the pJM17 adenovirus plasmid. Adenovirus particles were precipitated from 293T cell lysates using polyethylene glycol, followed by CsCl gradient centrifugation and gel filtration chromatography. Cells were infected with adenovirus by adding purified adenovirus directly to the cell culture medium at a concentration of 4 x 10^8^ IU/mL. pACCMV empty vector and pACCMV *NT5E* vector were also used as plasmids for transient transfections.

### TCF-LEF luciferase reporter assay

HEC-1-A and Ishikawa cells were seeded in 96-well plates at a density of 2.8 x 10^4^ cells/well. After 24 hours, cells were co-transfected with 100 ng TOPFlash/FOPFlash and 100 pg CMV-pRenilla-LUC, along with 100 ng wild type β-catenin, mutant β-catenin, or empty vector. All vectors were transfected into cells using Lipofectamine 3000 (Invitrogen) as per manufacturer instructions. At 48 hours post-transfection, cells were lysed with 100 ul passive lysis buffer (Biotium) and the luciferase assay was performed with the Firefly and Renilla Luciferase Single Tube Assay Kit (Biotium). Cell lysates were diluted 1:20 in 1X PBS for Firefly luciferase measurement and then 1:20 in 1X PBS again for Renilla luciferase measurement. Luciferase activity was measured using a TD-20/20 Luminometer (Turner BioSystems). *n* = 4-9 technical replicates were performed per condition for each experiment. Each dot represents one technical replicate, and data are reported as fold change vs. endogenous luciferase activity, unless otherwise stated.

### Immunofluorescence

Cells grown on 18×18 mm coverslips were fixed with 4% paraformaldehyde, incubated with 0.1% Triton X-100 and non-specific binding blocked with Background Snipper (Biocare Medical). Primary antibodies were incubated overnight at 4°C followed by appropriate secondary antibodies. Spectral bleed-through controls included the incubation of primary antibodies and related fluorochromes separately. For assessing nuclear localization of *Xenopus* myc-tagged exon 3 mutant β-catenin, ∼90 images at 20X magnification for each group (non-targeting or CD73 1247 siRNA) were captured from slides. Relative intensity for each image was measured using Keyence macro cell counting software.

### Immunohistochemistry in human tumors

Formalin-fixed paraffin-embedded tumor sections (4 μm) from *n* = 11 patient tumors validated by next generation sequencing to have exon 3 *CTNNB1* mutations were processed for immunohistochemistry for CD73 and β-catenin as previously described^17,32^. Tumor images (10-30 per sample, 10x magnification) were captured using a BX41 Olympus microscope. CD73 and β-catenin staining was manually quantified across *n* = 15-30 20x images per tumor section using cellSens software (Olympus).

### Real-time quantitative PCR

Quantitative RT-PCR for *NT5E* was performed on *n* = 32 endometrial cancer tissues validated to have exon 3 *CTNNB1* mutation by next generation sequencing. RNA was isolated from frozen tissues using TRIzol Reagent (Invitrogen), followed by purification with RNeasy columns (QIAGEN). Real-time qPCR was performed as previously described^17,31^.

### RNA extraction for RNA-sequencing

Cells were plated in 6-well plates at 5 x 10^5^ cells/well and then transfected at 85% confluency with empty vector (LentiV_Neo), D32N mutated β-catenin vector, G34R mutated β-catenin vector, or S37F mutated β-catenin vector (2 ug). At 48 hours post-transfection, cells were washed twice with 1X PBS, and the plates were placed at −80C. Total RNA isolation for *NT5E* WT (EV-, D32N-, G34R-, S37F-transfected) and *NT5E* KO (EV-, D32N-, G34R-, S37F-transfected) HEC-1-A cells was performed using QIAshredder and RNAeasy Mini Kits (QIAGEN) following manufacturer instructions. RNA concentrations were determined (Nanodrop) and samples were run on a 1% agarose gel to test integrity of 28 S and 18 S bands. Secondary assessment of quality controls and RNA-sequencing of the samples were performed by Novogene.

### RNA-seq analyses

Stranded Bulk mRNA sequencing was performed on Poly-T selected total RNA isolates following size distribution detection (Novogene). Transcriptomic sequences were gathered at 150bp paired-end reads at or exceeding 35 million reads per replicate. Several analyses were performed on the sequences: mapping by hisat2 (2.05), assembly by Stringtie (1.3.3b), quantification by featureCount (1.5.0-p3), and DE analysis by DESeq2 (1.20.0). R2 Genomics Analysis and Visualization Platform^120^ (http://r2.amc.nl) was used to generate heatmaps depicting log2 z-score for our literature search list of canonical Wnt/β-catenin pathway target genes and the β-catenin dependent gene list by Doumpas et. al.

### Western blot analysis

Total protein was isolated from cell extracts by either scraping frozen plates with RIPA buffer containing protease and phosphatase inhibitors (Thermo Fisher) or by lysing cells with the same buffer that were collected by trypsinization. Protein was quantified using protein assay dye (BioRad) and concentration measured at 595 nm. Equal amounts of protein (10 or 20 ug) per sample were separated by SDS-PAGE. Total protein was imaged using 2,2,2-trichloroethanol as previously described^121^. Immunoblotting was performed using PVDF (BioRad) or nitrocellulose membranes (BioRad). Proteins transferred to membranes were blocked in 5% (w/v) nonfat dry milk in 1X PBS containing 0.1% (v/v) Tween 20, incubated with primary antibodies at 4°C overnight, and expression detected by peroxidase-conjugated secondary antibodies and ECL chemiluminescence (Super Signal West Pico (Thermo Scientific)).

### Cellular fractionation

*NT5E* WT and *NT5E* KO HEC-1-A cells were plated in 150 cm plates at 6 x 10^6^ cells/plate and then transfected at 85% confluency with *Xenopus* β-catenin^ΔEX3^, G34R mutated β-catenin vector, or S37F mutated β-catenin vector (2 ug). At 48 hours post-transfection, cells were trypsinized, counted, and 12 x 10^6^ cells pelleted for cellular fractionation. A second pellet of 2 x 10^6^ cells was collected for total cell protein extract. Cellular fractionation was performed using a Pierce Subcellular Protein Fractionation Kit (Thermo Fisher). Total cell protein extracts were prepared using 1X RIPA buffer.

### Co-immunoprecipitation (Co-IP)

HEC-1-A *NT5E* WT and *NT5E* KO cells were transfected with *Xenopus* β-catenin^ΔEX3^, G34R mutated β-catenin vector, or S37F mutated β-catenin vector (2 ug). Co-IP was performed using a Pierce c-Myc-tag Magnetic IP/Co-IP Kit (Thermo Scientific). Protein (500 µg) was incubated overnight at 4°C with 25 μl anti-Myc magnetic beads. The resulting immune-bound complexes were eluted in 2X reducing sample buffer and assessed by SDS-PAGE and immunoblotting methods.

### Quantification and Statistical Analysis

P values were calculated using an unpaired t test, one-way ANOVA with Tukey/s post test, two-way ANOVA with Sidak’s post test, or as otherwise indicated (GraphPad Prism 10; GraphPad Software). Human tissue data were analyzed using a Mann-Whitney U test or Kruskal-Wallis one-way ANOVA with Dunn’s post test. A p value of less than 0.05 was considered significant. Survival data were collected by review of electronic medical records, and overall survival rates were stratified in a Kaplan-Meier plot according to *NT5E* mRNA levels. Densitometry on western blot images was performed using ImageJ^122^. Meta-Chart Venn Diagram Maker Online was used to make select Venn diagrams. KEGG database was used for RNA-seq analysis^123^.

## Conflicts of Interest

The authors declare no conflict of interest.

## Author Contributions

RMH and JLB developed the study design, performed experiments, and analyzed and interpreted the data. LFC and CV developed the *NT5E* adenoviral vector. SP and HNL performed immunoblots. KCK and RRB provided patient tissue samples and clinical data. EMR generated HEC-1-A CD73 control samples. RMH and JLB – writing of original draft of manuscript. RMH, SP, KCK, LFC, EMR, RRB, CV, and JLB – reviewing and editing of manuscript. Supervision – RRB, CV, JLB. Funding acquisition – RRB and JLB.

## Acknowledgements

We thank Pierre McCrea (University of Texas MD Anderson Cancer Center) for providing the *Xenopus* β-catenin^ΔEX3^ plasmid. We thank Gaith N. Droby for assisting with RNA-seq data management and methods preparation. We thank the R2 support team for work with the R2 Genomic Analysis & Visualization Platform. This work was supported by NIH P50CA098258 (RRB), a Career Enhancement Award (JLB) from NIH P50CA098258, and a University of North Carolina Center for Environmental Health and Susceptibility Pilot Award (JLB). JLB received support from an International Anesthesia Research Society Mentored Research Award. RMH and EMR were supported in part by the University of North Carolina Chapel Hill Program in Translational Medicine (NIH T32GM122741). LFC was supported in part by a grant from the National Institute of General Medical Sciences (5T32GM135128).

## Notes

### Competing Interest Statement

The authors have declared no competing interest.

