## Supplemental Material for "CD73 restrains mutant β-catenin oncogenic activity in endometrial carcinomas"

Supplemental Table 1

Clinicopathological features of qRT-PCR cohort of exon 3 *CTNNB1*-mutant endometrial carcinomas.

| **Characteristics (n = 29)** | **No Recurrence** | **Recurrence** |
|---|---|---|
| Histology | **n =** | **n =** |
| Endometrioid | 17 | 10 |
| Non-endometrioid | 2 | 0 |
| FIGO Stage | **n =** | **n =** |
| I | 14 | 7 |
| II | 1 | 0 |
| III | 3 | 2 |
| IV | 0 | 1 |
| Unknown | 1 | |
| Grade | **n =** | **n =** |
| G1 | 3 | 1 |
| G2 | 14 | 7 |
| G3 | 2 | 2 |
| Lymphovascular Space Invasion (LVSI | **n =** | **n =** |
| Yes | 5 | 5 |
| No | 14 | 3 |
| Unknown | | 2 |

Supplemental Table 2

Antibodies

| **Antibody** | **Source** | **Company** | **Catalog #/Clone** |
|---|---|---|---|
| Alexa Fluor 594 goat anti-mouse IgG | | Invitrogen | #A21206 |
| Alexa Fluor 594 goat anti-mouse IgG | | Invitrogen | #A11032 |
| β-catenin | Rabbit polyclonal | Cell Signaling | #8480/D10AB |
| CD73 | Rabbit | Cell Signaling | # 13160/D7F9A |
| E-cadherin | Mouse monoclonal | BD Biosciences | #36/E-Cadherin |
| GAPDH | Rabbit polyclonal | Cell Signaling | # 3683/14C10 |
| HRP-conjugated anti-mouse IgG | | Cell Signaling | #7076 |
| HRP-conjugated anti-rabbit IgG | | Cell Signaling | #7074 |
| Myc-tag (WB) | Rabbit polyclonal | Cell Signaling | #2278/ 71D10 |
| Myc-tag (IF) | Mouse monoclonal | Cell Signaling | #2276/9B11 |
| SP 1 | Rabbit polyclonal | Cell Signaling | #9389/D4C3 |
| Rab11a | Rabbit polyclonal | ABClonal | #A3251 |
| H2AX | Rabbit polyclonal | Cell Signaling | #7631/D17A3 |
| GSK3β | Rabbit monoclonal | Cell Signaling | #9312/27C10 |
| pGSK3β S9 | Rabbit polyclonal | Cell Signaling | #9323S/5B3 |
| α-catenin | Rabbit monoclonal | Cell Signaling | #3240/23B2 |
| GAPDH-HRP | Rabbit monoclonal | Cell Signaling | #3683S/14c10 |
| SOX17 | Rabbit polyclonal | ABclonal | #A18858 |
| FOSL1 | Rabbit polyclonal | ABclonal | #A5372 |

Supplemental Table 3

β-catenin target genes, downregulated in HEC-1-A *NT5E* KO cells

| **Gene** | **p-value, KO vs WT** |
|---|---|
| *GINS3* | 0.016997 |
| *SOX9* | 0.324417 |
| *UHRF1* | 0.043521 |
| *FOXRED2* | 0.056074 |
| *ABCC4* | 0.025321 |
| *SCARA3* | 0.037061 |
| *CCL28* | 0.00272 |
| *DKK1* | 0.011321 |
| *MNS1* | 0.002993 |
| *GINS2* | 0.05342 |
| *SLC7A2* | 0.015135 |
| *CLDN2* | 0.01134 |
| *SOX17* | 0.000201 |
| *MCM2* | 0.001181 |
| *ID2* | 0.000775 |
| *CCND1* | 0.002666 |
| *FAM111B* | 0.000191 |
| *RNF43* | 0.008008 |
| *ALDH1A1* | 0.000622 |
| *MMP7* | 0.006969 |

| | |
|---|---|
| *HDAC4* | 0.001222 |
| *EPHB3* | 0.002521 |
| *FZD7* | 0.000145 |

Supplemental Table 4

β-catenin target genes, upregulated in HEC-1-A *NT5E* KO cells

| Gene | p-value, KO vs WT |
|---|---|
| *ZNF367* | 0.147299 |
| *LGR5* | 0.101663 |
| *GJA1* | 0.24209 |
| *LOXL3* | 0.035689 |
| *PDK1* | 0.02015 |
| *CD3EAP* | 0.030654 |
| *PPARD* | 0.226127 |
| *NOS2* | 0.283438 |
| *SNAI1* | 0.016422 |
| *GBX2* | 0.040869 |
| *CD274* | 0.010798 |
| *TEAD4* | 0.063674 |
| *JUN* | 0.000082 |
| *PLAUR* | 0.005815 |
| *FOSL1* | 0.01345 |
| *DTL* | 0.005541 |
| *TFAP4* | 0.000273 |
| *FAM216A* | 0.000051 |
| *VEGFA* | 0.009235 |
| *CLDN1* | 0.001855 |
| *ANKRD13B* | 0.015263 |
| *GRAMD1A* | 0.002139 |
| *MYC* | 0.035353 |
| *JAG1* | 0.003937 |
| *CDT1* | 0.001499 |
| *FN1* | 0.100296 |
| *PALD1* | 0.018427 |
| *BAMBI* | 0.001473 |
| *NES* | 0.03024 |
| *STRA6* | 0.057879 |
| *TIAM1* | 0.061368 |

Supplemental Table 5

TCF-dependent Wnt target genes[1], downregulated in HEC-1-A *NT5E* KO cells

| Gene | p-value, KO vs WT |
|---|---|
| *TPI1P2* | 0.135108 |
| *NMNAT3* | 0.290828 |
| *PYGM* | 0.236633 |
| *MSX2* | 0.018796 |
| *SLC7A8* | 0.064381 |
| *HIST1H2BN* | 0.427417 |
| *BAG1* | 0.179839 |
| *HIST1H2AC* | 0.070826 |
| *HBP1* | 0.029965 |
| *CACNA1D* | 0.006801 |
| *BCL11B* | 0.021994 |
| *COL7A1* | 0.027357 |
| *TSHZ1* | 0.020746 |
| *THRB* | 0.00252 |
| *PAX2* | 0.011465 |
| *NEDD9* | 0.001869 |
| *B3GNT3* | 0.000015 |
| *CCND1* | 0.002666 |
| *DKK1* | 0.011321 |
| *CCDC87* | 0.028848 |
| *ELMO3* | 0.736051 |
| *LRIT3* | 0.649253 |
| *FERMT3* | 0.022604 |
| *SNAI3* | 0.008167 |

Supplemental Table 6

TCF-dependent Wnt target genes[1], upregulated in HEC-1-A *NT5E* KO cells

| Gene | p-value, KO vs WT |
|---|---|
| *GPR83* | 0.993146 |
| *UNC5C* | 0.186194 |
| *CYP2D7* | 0.780491 |
| *MYOM2* | 0.241658 |
| *EYA1* | 0.027181 |
| *CCNG2* | 0.024325 |
| *ZFAND2A* | 0.162111 |
| *PIP5KL1* | 0.079339 |
| *ANKRD24* | 0.020076 |
| *MSX1* | 0.014937 |
| *ANKRA2* | 0.000756 |
| *SYTL5* | 0.189944 |
| *HRH1* | 0.026711 |
| *THSD7A* | 0.405609 |
| *CCBE1* | 0.937148 |
| *KLLN* | 0.019551 |
| *PRPH* | 0.044281 |
| *GRM4* | 0.012582 |
| *MATN1-AS1* | 0.029069 |
| *HOXA9* | 0.130986 |
| *C1QL1* | 0.00512 |
| *JAK3* | 0.005432 |
| *TLL1* | 0.001282 |
| *RGS4* | 0.029391 |
| *SLC7A5P1* | 0.021268 |
| *INSC* | 0.000029 |
| *PLXNA2* | 0.03917 |
| *DDX60* | 0.114797 |
| *ADGRA2* | 0.090646 |
| *DOCK11* | 0.004414 |
| *PLAT* | 0.028507 |
| *RASSF6* | 0.008413 |
| *SYTL2* | 0.002717 |
| *CREBRF* | 0.008829 |
| *RASGEF1B* | 0.00022 |
| *NAP1L3* | 0.014844 |
| *SLC9A9* | 0.001679 |
| *TEX19* | 0.019683 |

| | |
|---|---|
| *RNF19B* | 0.007087 |
| *CLDN1* | 0.001855 |
| *ADAMTS5* | 0.005591 |
| *CALCB* | 0.012077 |
| *NTMT1* | 0.143381 |
| *ZNF503* | 0.000269 |
| *RPL41* | 0.152306 |
| *SOCS2-AS1* | 0.045362 |
| *SCUBE1* | 0.126216 |

Supplemental Table 7

Novogene-defined differentially expressed genes with D32N, G34R, and S37F in *NT5E* KO cells vs. EV *NT5E* WT cells.

| **Gene** | **Description** | **Mutant(s)** | **Δ, KO vs WT** | **Involvement in cancer** | **References** |
|---|---|---|---|---|---|
| *LINC01389* | Antisense | S37F | down | upregulation is associated with gastric cancer; correlated with EMT | [2] |
| *AC008438.1* | Antisense | S37F | down | upregulation in some cancers | [3] |
| *AL080317.1* | Antisense | S37F<br>G34R | down | upregulated in colon cancer; HR > 1 for Wilms tumor | [4,5] |
| *AC092119.2* | lincRNA | S37F<br>D32N | down | upregulated in gastric cancer, ccRCC | [6] |
| *LINC00113* | lincRNA | S37F<br>D32N | down | upregulated in TNBC, correlating with poor prognosis; up in lung cancer; downregulated in esophageal cancer; down in ccRCC | [7–10] |
| *MIR3613* | miRNA | S37F | down | down in CRC; tumor suppressor in breast cancer; | [11,12] |
| *TUBA1A* | protein_coding | S37F<br>G34R | down | up in GC; up in GBM | [13,14] |
| *HSPA1A* | protein_coding | S37F<br>G34R<br>D32N | down | up in breast cancer; CRC | [15,16] |
| *U2AF1* | protein_coding | S37F<br>D32N | down | Mixed/Unknown | [17,18] |
| *ZNF112* | protein_coding | S37F | down | Unknown | |
| *AP001107.4* | Antisense | G34R | down | down in CRC; | [19,20] |
| *AC010331.1* | Antisense | G34R | down | favorable prognostic factor in bladder cancer + breast cancer | [21,22] |
| *U62317.2* | Antisense | G34R<br>D32N | down | down in bladder cancer; high levels correlate with high OS in basal-like breast cancer; | [23,24] |
| *USP46-AS1* | lincRNA | G34R | down | increased OS in ccRCC and glioma; decreased in HCC | [25,26] |

| | | | | | |
|---|---|---|---|---|---|
| *AC127024.5* | lincRNA | G34R | down | HR 0.3 in pancreatic cancer | [27,28] |
| *AC005392.2* | lincRNA | G34R | down | pro-angiogenic in CRC; poor OS in AML | [29,30] |
| *AC124312.2* | lincRNA | G34R | down | risk factor for bladder cancer; | [31] |
| *AC005332.3* | lincRNA | G34R<br>D32N | down | HR 0.7 in pancreatic cancer; pro-EMT in HCC | [32–34] |
| *MIR5587* | miRNA | G34R | down | Unknown | |
| *MIR6783* | miRNA | G34R | down | Mixed/Unknown | [35] |
| *AC131235.3* | Antisense | D32N | down | shorter survival in CSC; | [36] |
| *FP236383.1* | lincRNA | D32N | down | Unknown | |
| *MIR641* | miRNA | D32N | down | prohibited prolif/met but promoted apoptosis in lung cancer; tumor suppressor in lung cancer and cervical cancer | [37,38] |
| *MIR3682* | miRNA | D32N | down | migration and stemness in HCC, through PI3K/β-catenin; | [39] |
| *MIR3939* | miRNA | D32N | down | associated with RT sensitivity in cervical cancer; associated with response to sunitinib in ccRCC; | [40,41] |
| *HIST2H2AA4* | protein_coding | D32N | down | associated with T2DM + pancreatic cancer; associated with hypothermic response in breast cancer | [42,43] |
| *HIST2H2AA3* | protein_coding | D32N | down | down in HCC; associated with T2DM + pancreatic cancer; associated w/ hypothermic response in breast cancer; up in brain mets in breast cancer | [42–45] |
| *NUDT4B* | protein_coding | D32N | down | does not cause CRC cell prolif; excluded from pan-cancer study due to similar expression in tumor vs normal tissue; | [46,47] |
| *U2AF1L5* | protein_coding | D32N | down | poor OS in NPC; | [48] |
| *F8A3* | protein_coding | D32N | down | lower OS in neuroblastoma; | [49] |
| *HIST3H2BB* | protein_coding | D32N | down | hypermethylated in lung cancer; | [50] |
| *ETV7* | protein_coding | D32N | down | oncogene and inta-inflammatory in breast cancer; promotes CRC; doxorubicin resistance in breast cancer; poor OS in bladder cancer | [51–54] |
| *VPREB3* | protein_coding | D32N | down | expressed on tumor cells with C-MYC translocations | [55] |

**Figure S1**

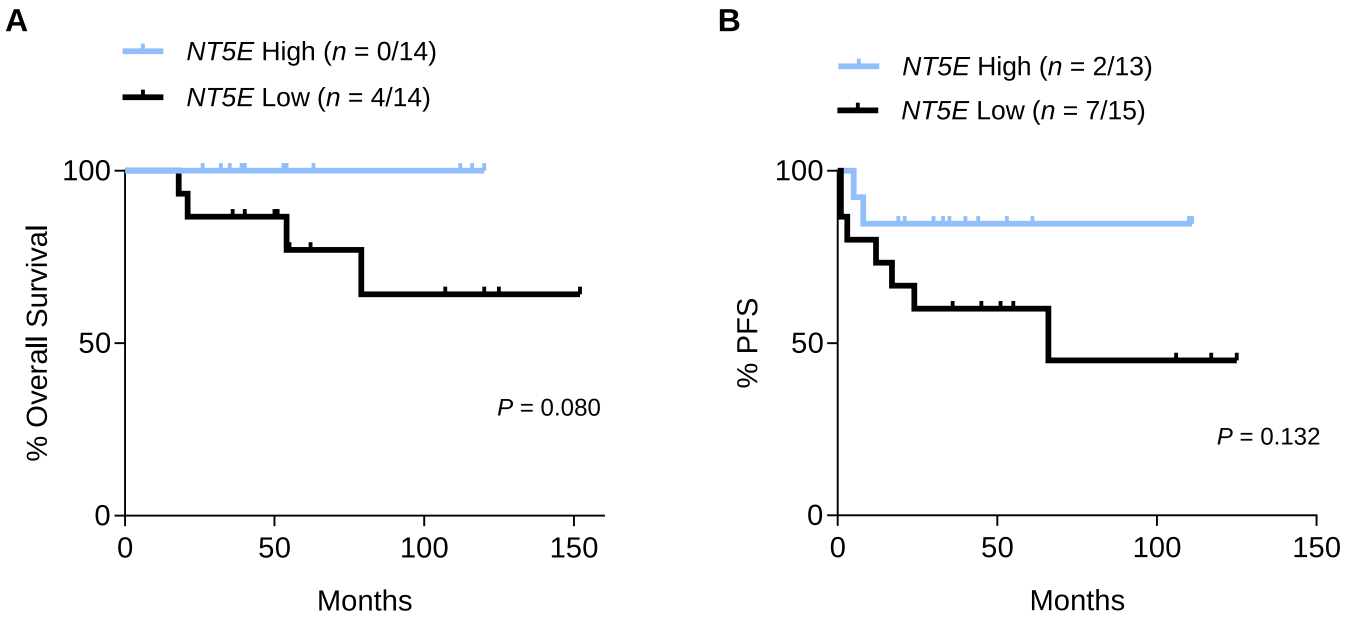
A
NT5E High (n = 0/14)
NT5E Low (n = 4/14)
100
50
0
% Overall Survival
P = 0.080
0
50
100
150
Months
B
NT5E High (n = 2/13)
NT5E Low (n = 7/15)
100
50
0
% PFS
P = 0.132
0
50
100
150
Months

#### Supplemental Figure Legends

**Supplemental Figure 1**. **Survival trends for patients with *NT5E* high and low exon 3 *CTNNB1* mutant tumors.** (**A**) Progression-free survival and (**B**) overall survival for patients with *NT5E* high and low endometrial tumors with exon 3 *CTNNB1* mutations. High and low values were determined by the median (0.001358). Logrank and Gehan-Breslow-Wilcoxon tests were used. $n$ = 28 patients for (**A**) and (**B**), survival data was missing for one patient from the $n$ = 29 cohort.

### Figure S2

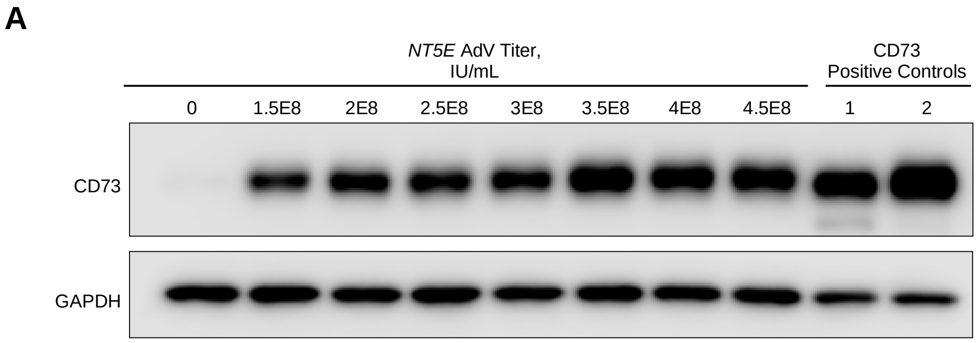

A
NT5E AdV Titer,
IU/mL
CD73
Positive Controls
0
1.5E8
2E8
2.5E8
3E8
3.5E8
4E8
4.5E8
1
2
CD73
GAPDH

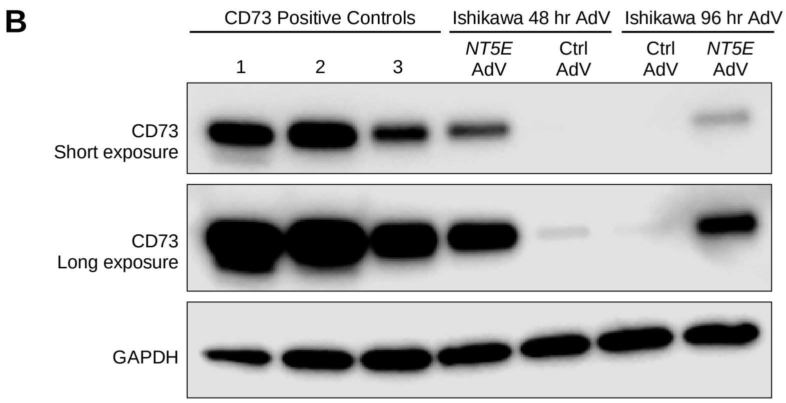

B
CD73 Positive Controls
Ishikawa 48 hr AdV
Ishikawa 96 hr AdV
1
2
3
NT5E AdV
Ctrl AdV
Ctrl AdV
NT5E AdV
CD73
Short exposure
CD73
Long exposure
GAPDH

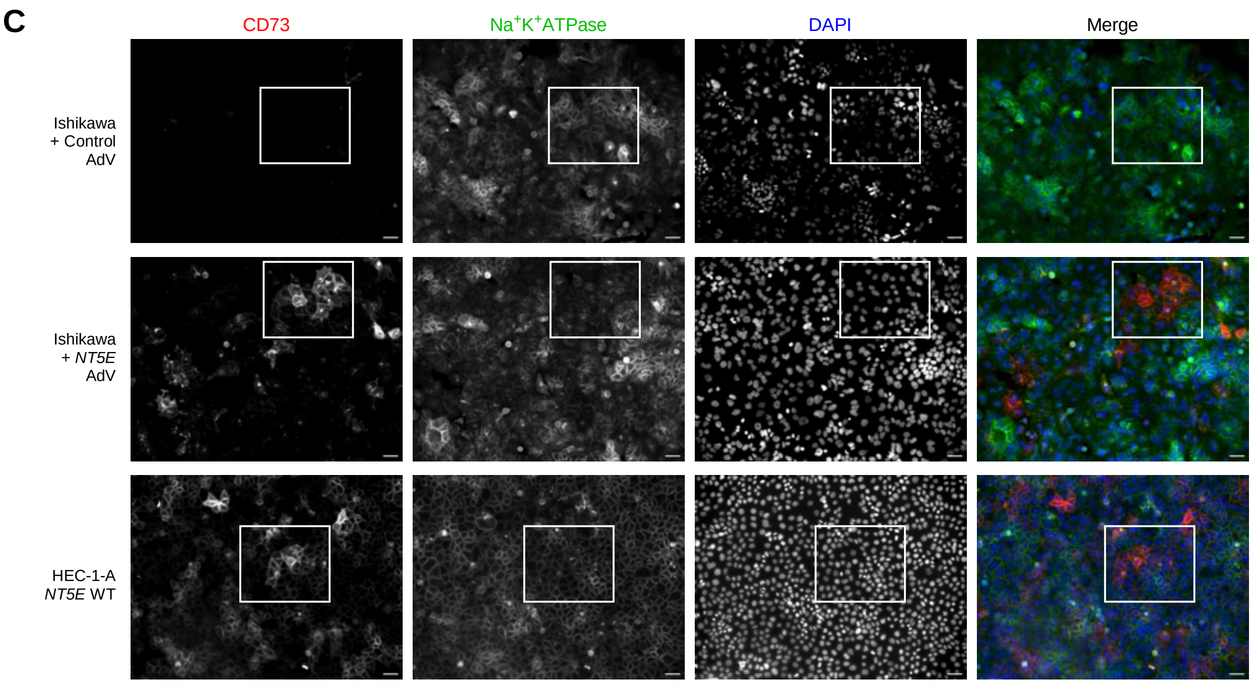

C
CD73
Na+K+ATPase
DAPI
Merge
Ishikawa
+ Control
AdV
Ishikawa
+ NT5E
AdV
HEC-1-A
NT5E WT

**Supplemental Figure 2. Induced expression of CD73 via *NT5E* adenovirus transduction in Ishikawa cells.** (**A**) CD73 protein expression with different *NT5E* AdV viral titers compared to HEC-1-A cells which serve as positive controls. CD73 Positive Control 1 = HEC-1-A cells at 100% confluency, 2 = HEC-1-A cells at 2 days post-confluency. (**B**) Validation of continued CD73 expression in Ishikawa cells. Expression persists for 96 hours, the endpoint in which TCF/LEF luciferase assays were performed. HEC-1-A cells serve as CD73 positive controls. Lanes 1 = HEC-1-A cells at 100% confluency; 2 = HEC-1-A cells at 2 days post-confluency; 3 = Ishikawa cells previously transduced with *NT5E* AdV and no luciferase reporter plasmids. NT = no transduction. (**C**) Uncropped immunofluorescence images corresponding to Fig. 2H. Cropped areas indicated with white rectangle. Scale bars 20 um.

### Figure S3

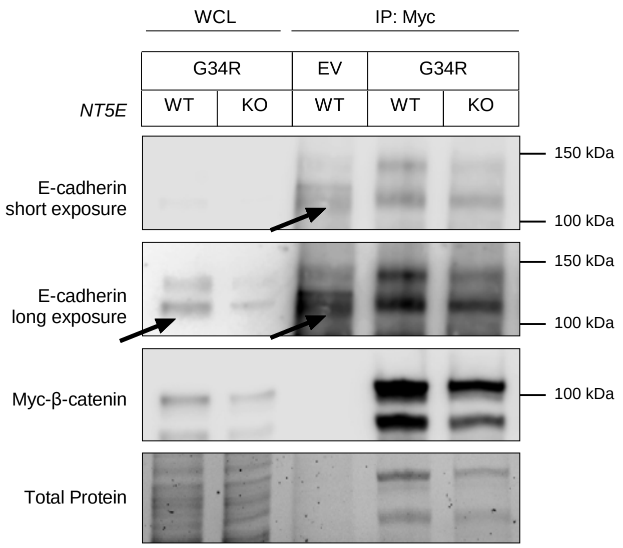

WCL
IP: Myc
G34R
EV
G34R
NT5E
WT
KO
WT
WT
KO
E-cadherin
short exposure
150 kDa
100 kDa
E-cadherin
long exposure
150 kDa
100 kDa
Myc-β-catenin
100 kDa
Total Protein

**Supplemental Figure 3. Patient-specific exon 3 β-catenin mutant binds with E-cadherin.** Immunoblots from co-immunoprecipitation experiment in *NT5E* WT and *NT5E* KO HEC-1-A cells. Myc-β-catenin was precipitated, and samples were probed for E-cadherin and myc-β-catenin expression G34R. *NT5E* KO cells have decreased expression of E-cadherin compared to *NT5E* WT cells. Thus, *NT5E* KO cells have a reduced capacity for E-cadherin-β-catenin binding. Accordingly, *NT5E* KO cells have weaker cell adhesions and increased nuclear β-catenin.

### Figure S4

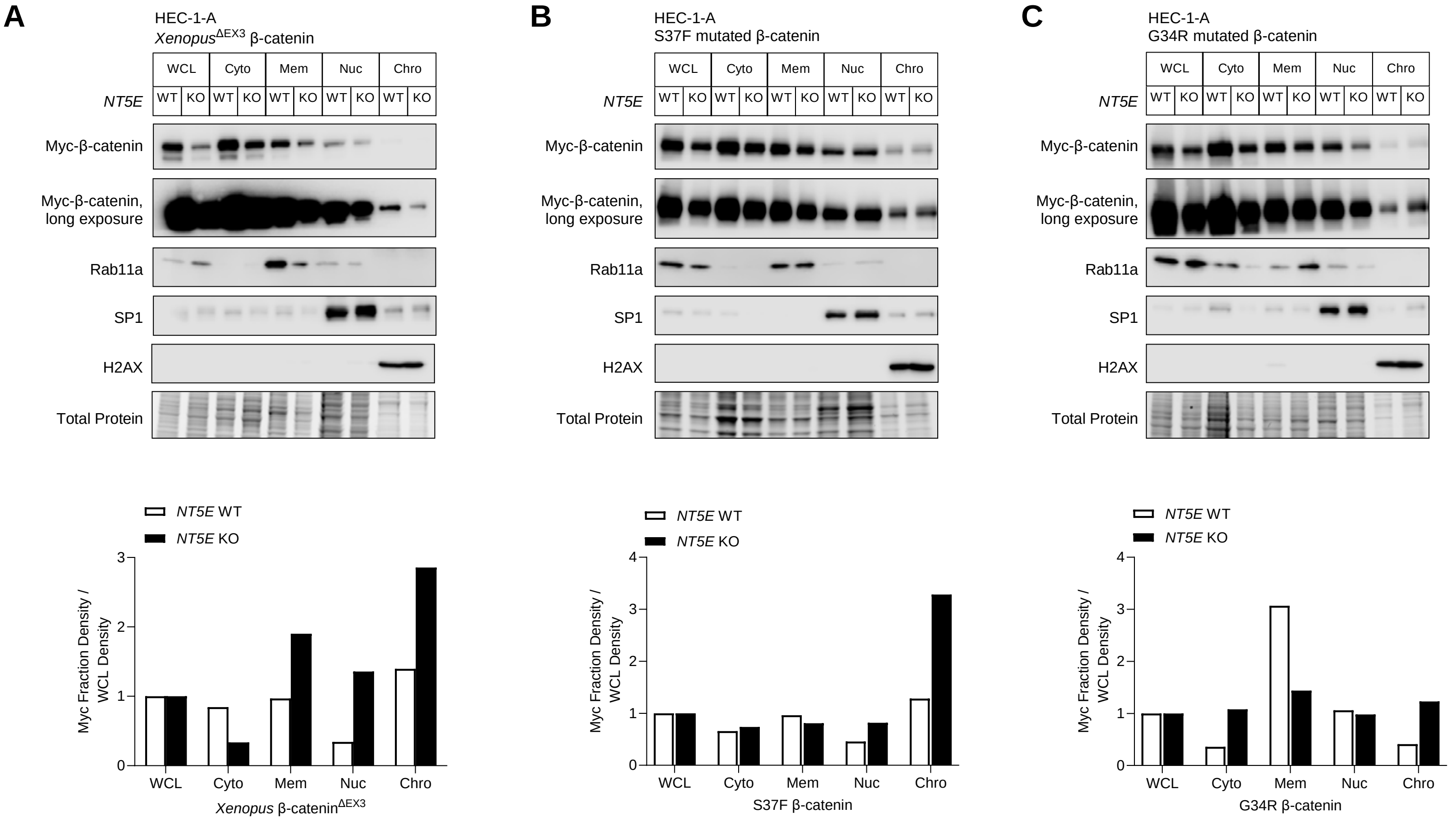

A
HEC-1-A
Xenopus ΔEX3 β-catenin
B
HEC-1-A
S37F mutated β-catenin
C
HEC-1-A
G34R mutated β-catenin
WCL
Cyto
Mem
Nuc
Chro
NT5E
WT
KO
Myc-β-catenin
Myc-β-catenin, long exposure
Rab11a
SP1
H2AX
Total Protein
NT5E WT
NT5E KO
Myc Fraction Density / WCL Density
Xenopus β-catenin ΔEX3
S37F β-catenin
G34R β-catenin

**Supplemental Figure 4. Independent replicates of cellular fractionations with patient-specific β-catenin mutations**. (**A**, **B, C**) Independent replicates of the cellular fractionation experiments described in Fig. 4D, 4E, and 4F from *NT5E* WT and *NT5E* KO HEC-1-A cells. *NT5E* WT and *NT5E* KO HEC-1-A cells were transfected with patient-specific β-catenin mutants (**A**) *Xenopus* β-catenin$^{\Delta EX3}$, (**B**) S37F, or (**C**) G34R. Densitometry graphs are shown for myc-β-catenin mutant expression for each cellular fraction normalized to myc-β-catenin mutant expression in the whole cell lysate (WCL). Cellular fraction markers: Rab11a (membrane), SP1 (nuclear), and H2AX (chromatin).

### Figure S5

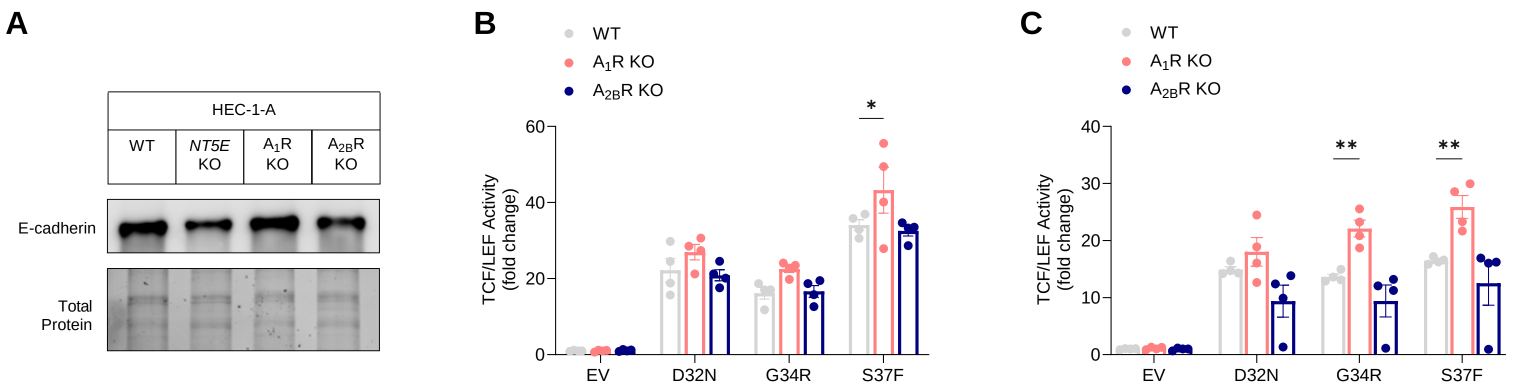
A
HEC-1-A
WT
NT5E KO
A1R KO
A2BR KO
E-cadherin
Total Protein
B
WT
A1R KO
A2BR KO
*
TCF/LEF Activity (fold change)
60
40
20
0
EV
D32N
G34R
S37F
C
WT
A1R KO
A2BR KO
**
**
TCF/LEF Activity (fold change)
40
30
20
10
0
EV
D32N
G34R
S37F

**Supplemental Figure 5. Reduced transcriptional activity of patient-specific β-catenin mutants in $A_1$R KO cells.** (**A**) Immunoblot showing E-cadherin expression is unchanged in HEC-1-A WT, *ADORA1* KO, and *ADORA2B* KO cells, but decreased in *NT5E* KO cells. (**B-C**) Independent replicates for data shown in Figure 6D. TCF/LEF reporter activity in cells transfected with empty vector (EV) or patient-specific β-catenin mutants D32N, G34R, or S37F. Each dot represents one technical replicate. Data represent mean ± SEM. *P < 0.05, **P < 0.01; 2-way ANOVA with Dunnett's post test.

### Figure S6

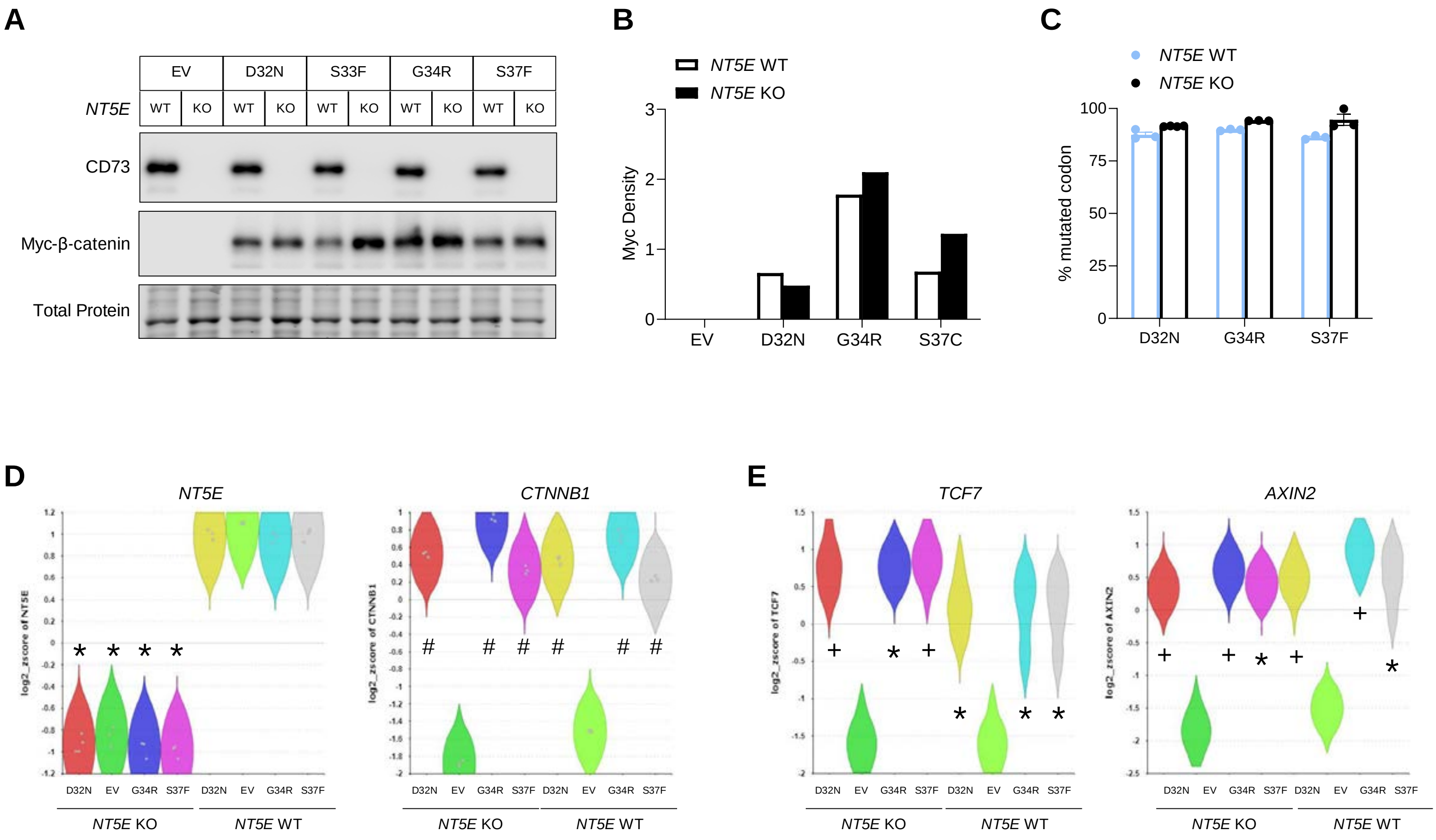

*P < 0.001 vs. *NT5E* WT
$^{\#}$P < 0.0005 vs EV

*P < 0.05 vs. EV
$^{+}$P < 0.005 vs. EV

**Supplemental Figure 6. Validation of patient-specific β-catenin mutant expression and activity in RNA-seq samples.** (**A**) Protein samples were collected in sync with samples process and submitted for RNA-sequencing. Immunoblots were used to assess equal or near equal expression of each patient-specific β-catenin mutant between *NT5E* WT and *NT5E* KO HEC-1-A cells. Due to unequal protein expression of S33F between *NT5E* WT and *NT5E* KO samples, RNA from these samples was not submitted for sequencing. **(B**) Densitometry for myc-β-catenin from samples in (**A**) used for RNA-sequencing. (**C**) Mutation frequencies for D32N, G34R, and S37F, calculated using Integrative Genomics Viewer[56]. (**D-E**) Validation of our experimental system using mRNA expression levels from RNA-seq data. (**D**) As expected, NT*5E* levels were low in *NT5E* KO samples and *CTNNB1* expression increased in both *NT5E* KO and *NT5E* WT samples expressing patient-specific β-catenin mutants. (**E**) Validation of β-catenin mutants to induce Wnt/β-catenin signaling gene targets, *TCF7* and *AXIN2*, is shown.

### Figure S7

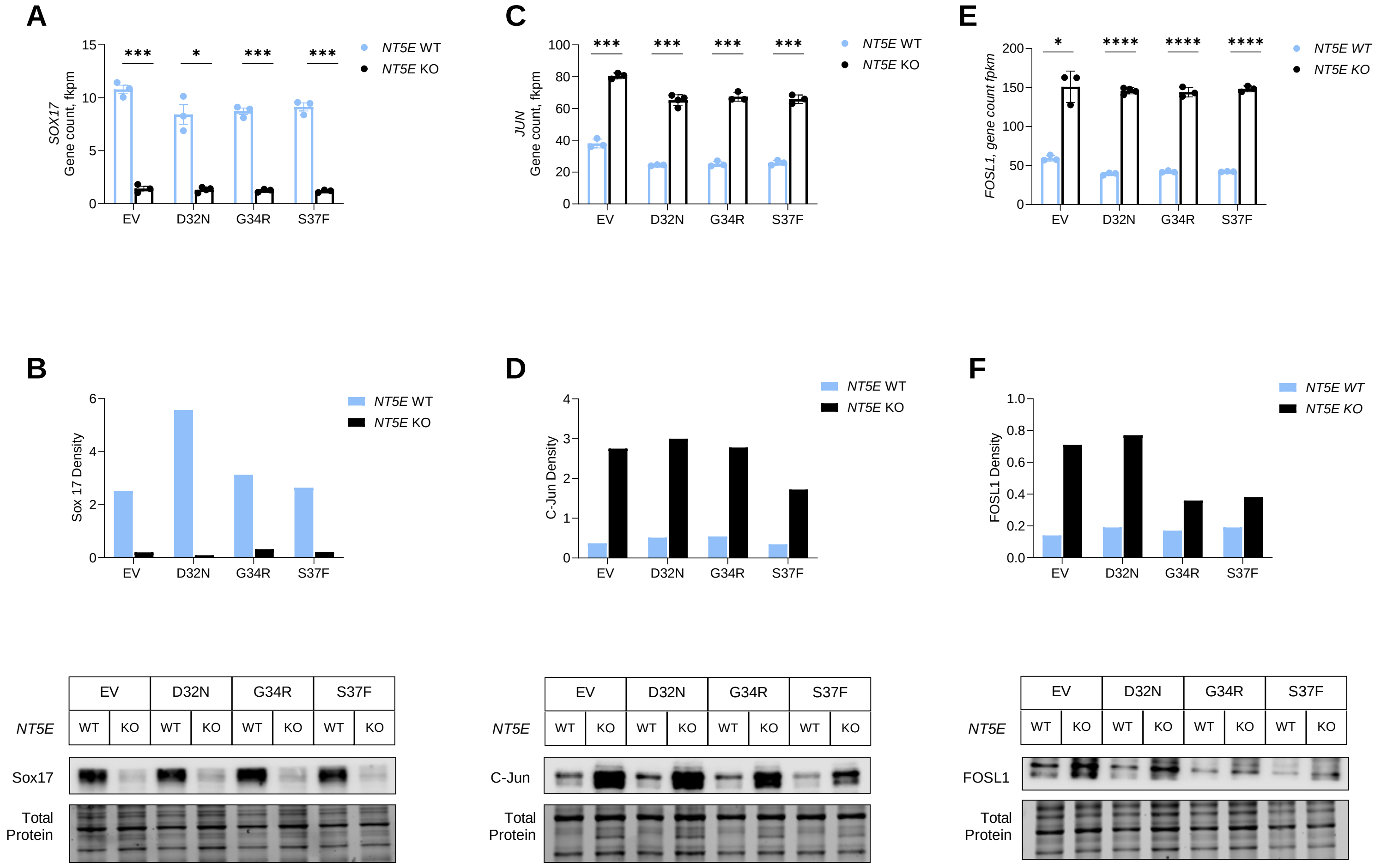

A
SOX17 Gene count, fkpm
NT5E WT
NT5E KO
EV
D32N
G34R
S37F
***
*
***
***
C
JUN Gene count, fkpm
***
***
***
***
E
FOSL1, gene count fpkm
*
****
****
****
B
Sox 17 Density
D
C-Jun Density
F
FOSL1 Density
NT5E
WT
KO
Sox17
C-Jun
FOSL1
Total Protein

**Supplemental Figure 7. Validation of dysregulated genes in *NT5E* WT and *NT5E* KO HEC-1-A cells identified from RNA-seq studies.** (**A**) mRNA expression and (**B**) protein expression of Sox17 in *NT5E* WT and *NT5E* KO cells. (**C**) mRNA expression and (**D**) protein expression of C-Jun in *NT5E* WT and KO cells. (**E**) mRNA expression and (**F**) protein expression of Fra1 in *NT5E* WT and KO cells. (**B**, **D, F**) Representative immunoblots for *n* = 3 independent experiments. *P < 0.05, **P < 0.01, ***P < 0.005, Welch t-test.
